# Preliminary pharmacokinetics and *in vivo* studies indicate analgesic and stress mitigation effects of a novel NMDA receptor modulator

**DOI:** 10.1101/2024.06.22.600208

**Authors:** Blaise M. Costa, De’Yana Hines, Nakia Phillip, Seth C. Boehringer, Ramu Anandakrishnan, McAlister Council-Troche, Jennifer L. Davis

**Author notes:** Blaise M. Costa, PhD, Edward Via Virginia College of Osteopathic Medicine Center for One Health Research, Virginia Tech, 1410 Prices Fork Rd, Blacksburg, VA 24060. **List of nonstandard abbreviations:** Cmax: maximum concentration; CNS4: Costa NMDAR stimulator 4; CS: Conditioned stimuli; FC: Fear condition; GluN: Glutamate receptor NMDA subtype; LOQ: Limit of quantification; NMDAR: N-methyl D-aspartate receptor; Tmax: time taken to reach maximum concentration; US: Unconditioned stimuli. UPLC-MS: Ultra-performance liquid chromatography-mass spectrometry.

## Abstract

NMDAR channel blockers produce analgesic and antidepressant effects by preferentially inhibiting the GluN2D subtype at lower doses. Given the distinct physiological role of GluN2 subunits, we hypothesized that compounds capable of simultaneously modulating GluN2A and GluN2D subtypes in opposite directions could serve as effective analgesics with minimal cognitive adverse effects. In this translational study, we investigated the *in vivo* effects of CNS4, a recently discovered glutamate concentration-dependent NMDAR modulator. Pharmacokinetic data revealed that CNS4 reaches peak plasma and brain concentrations within 0.25 hours post-intraperitoneal injection, with brain concentrations reach values up to 8.4% of those in plasma (64.9 vs. 5.47 µg/ml). Preliminary results showed that CNS4, a non-opioid compound, increased escape latency in mice during a hotplate assay by 1.74-fold compared to saline. In a fear conditioning (FC) experiment, CNS4 anecdotally reduced the electric shock sensation and significantly decreased stress-related defecation (fecal pellets: males, 21 vs. 1; females, 19 vs. 3). CNS4 also improved hyperarousal behavior (25 vs. 4 jumps), without affecting fear memory parameters such as freezing episodes, duration, or latency. CNS4 caused no changes in locomotion across 8 of 9 parameters studied. Remarkably, approximately 50 hours after FC training, CNS4 prevented stress-induced excessive sucrose drinking behavior by more than 2-fold both in male and female mice. These findings suggest that CNS4 penetrates brain tissue and produces pharmacological effects like those of NMDAR-targeting drugs but with a distinct mechanism, avoiding the undesirable side effects typical of traditional NMDAR blockers. Therefore, CNS4 holds potential as a novel non-opioid analgesic, warranting further investigation.

**Significance:** NMDA-subtype glutamate receptors are an attractive target for chronic pain and PTSD treatments as they play a critical role in forming emotional memories of stressful events. In this translational pharmacology work, we demonstrate the central analgesic and stress-mitigating characteristics of a novel glutamate concentration-biased NMDA receptor modulator, CNS4.

**Visual Abstract:** 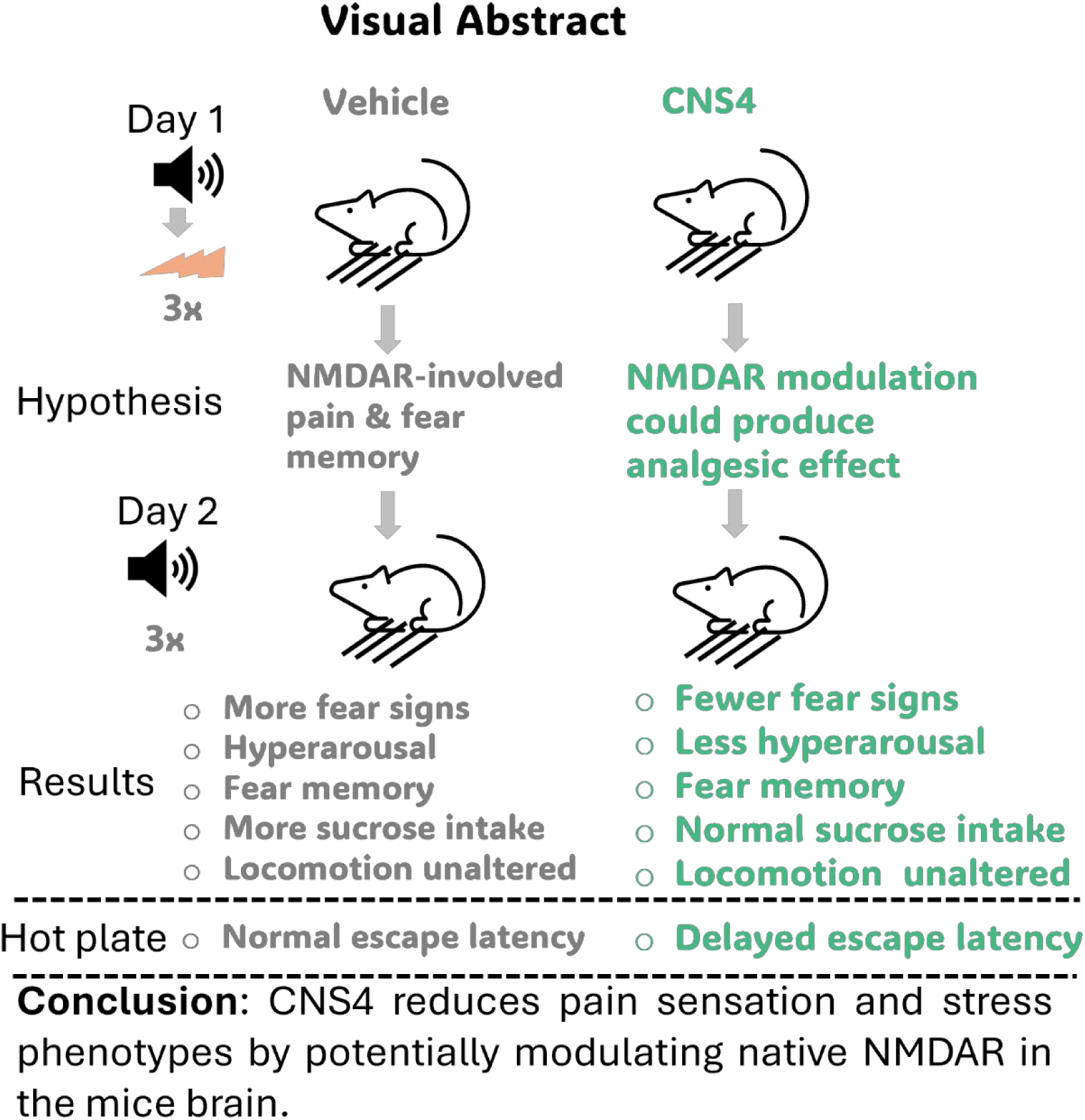

## 1. Introduction

Functional glutamatergic N-methyl-D-aspartate receptors (NMDARs) consist of two identical glycine-binding GluN1 subunits and two non-GluN1 subunits, which can be either similar or different. There are four glutamate binding GluN2 subunits (GluN2A, GluN2B, GluN2C, and GluN2D) that form functional di- or tri-heteromeric NMDARs (Monyer et al., 1992). Two GluN3 (GluN3A & 3B) subunits also have been identified to form excitatory glycine receptors by co-assembling with GluN1(Chatterton et al., 2002). Given their widespread physiological role, dysfunction of NMDARs causes various neurological and psychiatric conditions (Hansen et al., 2017); one of the complicated conditions is pain. Pain affects more people than diabetes, heart disease, and cancer combined, and is the most common reason people seek medical attention (Wager, 2022). Pain restricts daily life activities and the treatment of pain with opioids has contributed significantly to the opioid addiction epidemic and depression (Dahlhamer et al., 2018; Gureje et al., 1998; Smith et al., 2001). A balanced interplay between the excitatory glutamatergic neurons and GABAergic inhibitory neurons is essential to initiating and maintaining central pain sensitization (Breitinger and Breitinger, 2023). NMDARs play a pivotal role in central pain sensitization, which evolves from activation of neuronal circuits in nociceptive pathways (Aiyer et al., 2018; De Kock and Lavand’homme, 2007; Petrenko et al., 2003). Chronic pain complaints are common in patients with a primary diagnosis of post-traumatic stress disorder (PTSD), with prevalence rates estimated to be as high as 80% (Lew et al., 2009; Otis et al., 2009). Both PTSD and chronic pain involve alterations in brain regions related to emotion, cognition, and sensory processing (Burris et al., 2009; Jenewein et al., 2009; Sharp and Harvey, 2001), where NMDARs are expressed. Therefore, these two pathogenically connected and co-existing conditions could be altered by pharmacological agents that modulate NMDAR function.

NMDAR serves as a major non-opioid analgesic drug target (Ghoreishi-Haack et al., 2018; Houck et al., 2019). Particularly, compounds that modulate the GluN2D subtype could be useful in treating neuropathic pain (Dedek and Hildebrand, 2022; Hildebrand et al., 2014; Jing et al., 2022). The potential utility of such compounds derives from two unique characteristics of GluN2D: (1) GluN2D expression in adults is largely confined to regions that are involved in neuropathic pain perception, such as the thalamus, basal ganglia, and GABAergic interneurons across the brain and spinal cord (Dubois et al., 2016; Logan et al., 2007; Standaert et al., 1996; Tolle et al., 1993; Wenzel et al., 1996); (2) As the GluN1/2D channel is the slowest NMDAR subtype to deactivate, it can conduct a large number of cations into the neurons and trigger action potentials (Chen et al., 1999; Cull-Candy and Leszkiewicz, 2004; Erreger et al., 2005). Together, GluN2A and GluN2D subtypes of NMDARs play a critical role in excitatory and feed-forward inhibitory neurotransmission, as they are predominantly expressed on primary neurons and GABAergic interneurons, respectively (Dubois et al., 2016; Hansen et al., 2021; Logan et al., 2007; Mohrmann et al., 2000; Petralia et al., 2010; Standaert et al., 1996; Thomas et al., 2006; Tolle et al., 1993; Wenzel et al., 1996). Of note, a highly effective, dose-dependent, anti-depressant activity of esketamine is believed to be mediated through inhibition of the GluN2D subtype of NMDARs (Han et al., 2021; Hanson et al., 2024; Ide et al., 2017), demonstrating that direct inhibition and indirect disinhibition of primary neurons through GluN2D receptors have immediate and profound clinical effects (Han et al., 2021; Hanson et al., 2024; Ide et al., 2017; Kavalali and Monteggia, 2012).

Therefore, we hypothesize that pain sensation-mediated glutamatergic dysfunction could be corrected by concomitantly modulating GluN2A and GluN2D subunits in opposite directions. This hypothesis fits with the pharmacology of a recently discovered glutamate dose-dependent NMDAR modulator, 4-fluoro-N-[[2-(3-pyridinyl)-1-piperidinyl]-sulfanylidenemethyl]benzamide, CNS4 (Boehringer et al., 2023; Costa, 2018; Costa et al., 2021). As described previously (Costa et al., 2021), this compound preferentially potentiates neuroprotective GluN2A receptors by 22% and inhibits 35% of GluN2D when activated by 300µM glutamate. Conversely, CNS4 potentiates GluN2D currents by 8-fold when activated by 0.3µM glutamate. Similar glutamate dose-dependent activities of CNS4 on other GluN2 subtypes have recently been reported (Boehringer et al., 2023; Costa, 2018; Costa et al., 2021). In this work, we have initially performed preliminary pharmacokinetics and *in vivo* characterization to study the analgesic effect of CNS4. Thereafter, we have continued to study the effect of CNS4 on fear memory using an electric shock-induced fear conditioning (FC) model. Overall, pharmacokinetic profiling and *in vivo* assessments reveal central analgesic and stress-mitigating activities of CNS4.

## 2. Methods

### Animals

All animal experimental procedures and housing were approved by the Institutional Animal Care and Use Committee (IACUC) of Virginia Tech, protocol number 23-151. Thirty adult male CF-1 mice were used for pharmacokinetics work. Forty C57BL/6J mice (20 females and 20 males) were used for analgesic, FC, open field, and sucrose preference assay. Experiments were carried out in two phases. Twenty mice (10 males and 10 females) were studied in the first phase. These animals were procured from Charles River Laboratories (CRL). In the second phase, experiments were carried out on twenty mice (10 males and 10 females) procured from Jackson laboratories (JAX). We acknowledge that inter-strain differences in pharmacokinetics might exist, which could affect any pharmacokinetic and/or pharmacodynamic relationships.

Compound: CNS4 was synthesized through a contract research organization, Syngene International Limited, Bangalore, India. CNS4 used in this study met pharmaceutical-grade standards with >98% purity as verified by HPLC, NMR and LCMS assays, and as previously published (Boehringer et al., 2023; Costa et al., 2021).

### Pharmacokinetics (PK)

CNS4 was initially dissolved in 100% DMSO (Sigma-Aldrich, USA, cat#D-8779), then diluted in an appropriate volume of sterile normal saline and one equivalent of sodium hydroxide to get a fully solubilized clear solution. The final concentration of DMSO was 5%. CF-1 male mice were injected intraperitoneally with 100mg/kg CNS4 using a 27-gauge needle. Injection volume was between 100-200µl based on animal body weight. Animals were euthanized at 0, 0.25, 0.5, 1, 2, 4, 8, 12, 24, and 48 hours after CNS4 administration to collect blood and brain tissue samples. Three animals were used for each time point. Blood samples were collected in 0.4ml heparin EDTA vials and gently mixed before centrifuging at 2500 x g for 15 minutes to separate the plasma. The whole brain was excised and placed into vials containing RNA later solution (Thermofisher, USA, cat# AM7021) maintained at 4°C. One-half of the brain tissue samples were flash-frozen in liquid nitrogen and stored at −80°C. The other half was homogenized in 1x phosphate buffered saline and spun at 3200 x g for 30 minutes at 4°C and the supernatant was used for determination of drug concentration.

Plasma and brain samples were analyzed for the parent compound using UPLC-MS/MS. Plasma and brain standards and samples (20 μL) were mixed with 600 μL of methanol (MeOH) containing an internal standard solution using CNS43 and CNS41(Syngene International, India) at a concentration of 11.25 ng/mL. Standard curves were linear over a range of 0.003 – 1.5 μg/mL (R^2^ > 0.99). Quality control samples were analyzed in triplicate at concentrations of 0.003, 0.03, 0.3 and 1.5 μg/mL with a combined mean (± SD) intraday accuracy and precision of 100.2% (±0.74%) and 3.16% (± 4.26%), respectively. The limit of quantification (LOQ) was 0.003 µg/mL, and the limit of detection (LOD) was 0.00025 μg/mL based on a signal:noise ratio (S/N) of 3.

Chromatographic separation was performed on a Waters H-Class UPLC system with HSS T3 column (Waters Acquity UPLC HSS T3, 100 mm length x 2.1 mm ID x 1.7 µm) and matching guard column (Waters Acquity UPLC HSS T3 VanGuard Pre-Column, 5 mm length x 2.1 mm ID x 1.7 µm) maintained at 30°C. One microliter of sample was injected onto the column using a refrigerated autosampler maintained at 10°C. Mobile phase A consisted of 1% formic acid (FA) in water, and mobile phase B consisted of 0.5% FA + 2.5mM ammonium formate in methanol. The mobile phase was delivered to the UPLC column at a flow rate of 0.3 mL per min. The gradient elution program used was 80% A and 20% B for 0-3 minutes; 10% A and 90% B for 3-5 minutes; and 80% A and 20% B for 5-6 minutes. Retention time for the parent compound was approximately 1.7 min. Detection was performed on a triple-quadrupole mass spectrometer (Waters Xevo TQD) equipped with Zspray ionization, operated in positive-ion electrospray mode (ESI+) using multiple reaction monitoring (MRM). Commercial software (MassLynx) was used to analyze the data.

The extracted samples were screened for the presence of metabolites, as predicted using Swiss ADME software (Daina et al., 2017). The extracted samples were scanned for the presence of metabolites, as predicted using Swiss ADME software [142]. This program predicted CNS4 undergoing CYP450 3A4 mediated oxidation at the sulphonyl group (Metabolite-1) in the phase-1 metabolic reaction, and glucuronidation (Metabolite-2) at the nitrogen atom of the pyrimidine ring in the phase-2 reaction.

Pharmacokinetic data was then generated using naïve pooled concentrations and noncompartmental analysis for sparse sampling with Phoenix WinNonlin, version 8.4, Certara, Princeton, NJ.

### Hot plate assay

Three groups of adult C57BL mice were used for this experiment. About 15 minutes before the experiments, 7 mice (4 males and 3 females) were injected with sterile normal saline (0.9% NaCl), 6 mice (3 males and 3 females) with 100mg/kg CNS4 in normal saline, and 7 mice (3 males and 4 females) with 5mg/kg of meloxicam (Thermo Scientific, USA, cat# J60635.06) in normal saline, through intraperitoneal (IP) route. Ugo Basile Hot/Cold Plate equipment was used for this assay. The plate was digitally set to 55°C throughout the experiment. Although temperatures in the range of 45-50°C are generally used for this type of experiment, we used a higher temperature of 55°C to produce a pain stimulus that requires greater central nervous system integration and processing, making it appropriate for evaluating centrally-acting analgesics (Deuis et al., 2017; Hijazi et al., 2017). Mice were individually placed on the hot plate and quickly removed after they licked the paw for the first time. The time interval between placement and removal was noted as paw-licking (escape) latency. Statistical analysis of this data was performed using one-way ANOVA and post hoc Tukey’s multiple comparison tests. GraphPad Prism 10.2.3 software program was used for all statistical analysis. The same cohort of mice was used for the fear conditioning assay 5 days later.

### Open field assay (OFA)

A rodent open field apparatus (Maze engineers, part# 3203) was used for this assay. The AnyMaze 7.4 video recording system (Stoelting Co. IL) was used to monitor animal behavior, data collection, analysis, and statistics. Each mouse was individually placed in the apparatus for three minutes, and its uninterrupted movement was recorded and analyzed for nine different parameters presented in the results section. Freezing and immobility were calculated from 3 seconds cutoff. Four groups of mice were used for the OFA, sucrose preference test, and FC experiments (control male, control female, CNS4 male & CNS4 female). Each group contained 10 mice. All parameters were initially analyzed using two-way ANOVA, then post hoc Tukey’s multiple comparisons test, to find out differences between individual groups.

### Sucrose preference test (SPT)

Two identical 50ml tubes with drip nozzles were filled with an equal volume of water or 1% weight per volume of sucrose (Thermofisher, USA, cat# A15583.0E) in water solution. Regular water bottles were replaced, for about 16 hours (overnight), with tubes on the same side by inserting the nozzles into the cage where the original water bottle was present. Each tube was weighed before and after the experiment and the difference was calculated. Two sets of SPT were performed, before and after the FC experiment. Each of these groups had an equal number (five) of mice in each cage. Since mice were caged together as a group and received foot shock as a group, the SPT was performed as a group instead of per individual animal to avoid the influence of social isolation on fluid intake. Water and sucrose vials were placed on an empty cage to account for any inadvertent loss of volume due to evaporation or leakage. The fold changes in the sucrose-to-water preference ratio were used for the analysis.

### Fear conditioning (FC) assay

Maze Engineers’ (part# 5801) FC apparatus was used for this study. Conditioned stimuli (CS) and unconditioned stimuli (US) were precisely delivered into the chamber in a sequential manner through a software program, ConductMaze, that comes with the apparatus. The FC apparatus contains an acrylic cage with dimensions 17 x 17 x 25cm (width x depth x height), with four walls and no top and no base, a circular door that opens in the middle, and a floor grid that delivers foot shock. The apparatus is equipped with dual speakers on the backside inside of the chamber that deliver white noise at a tone of frequency 100-20,000 Hz, with a volume between 1-130dB. Before each test, the acrylic walls and floors were cleaned with water, while the shock grids were wiped with 70% ethanol to prevent interference of olfactory cues from previous animals. Mice were housed in their regularly enriched home cages before being transferred to a separate room where the FC apparatus was placed. All five mice in a group underwent the day 1 FC training protocol together and returned to their home cage before returning to their original housing room. This methodology is often employed to save time during the training phase (Chang et al., 2009; Shoji et al., 2014). However, mice were tested individually on day 2 for their fear response associated with FC. Below are the details on duration and intensity of CS and US: Tone, volume=50, duration=26s, frequency=4kHz; white noise, volume=50, duration=2s, 20kHz; shock, 0.5mA, 2s. On FC training day, CS (tone & white noise) was paired with US (foot shock); and on FC testing day, CS was unpaired with US, as there was just the sound (tone & white noise) but no shock.

About 15 minutes before the FC training, control male (Sal-M) and control female (Sal-F) groups were injected with sterile normal saline (0.9% NaCl), and CNS4 male (CNS4-M) and CNS4 female (CNS4-F) groups with 100mg/kg CNS4 in normal saline through intraperitoneal (IP) route. On testing day, animals underwent the CS (unpaired with US) protocol individually. Freezing episodes, duration, latency, hyperarousal (jumping behavior), and number of fecal pellets of individual animals were monitored during their time in the FC chamber, using the Any-Maze video recording program, and analyzed. Since animals were allowed to acclimatize in the chamber for the first three minutes (180 seconds), freezing data analysis was done for the 181-225s (tone 1), 227-271s (tone 2), and 272-316s (tone 3) of the stimulus protocol. These points cover 44s since they began hearing the tone. Freezing was calculated from 3 seconds immobility cutoff. Freezing data sets were initially analyzed using two-way ANOVA, then Šídák’s or Tukey’s multiple comparisons post hoc tests, to determine differences between individual groups. Hyperarousal (jumping behavior after placing the animal inside the FC chamber) was monitored throughout the duration inside the chamber on testing day.

A picture of the dirt collection tray, located underneath the FC chamber, was taken for each group (training day) or individual mouse (testing day) after completion of the stimulus protocol. The fecal pellets were counted and compared between saline and CNS4-treated groups using the Mann-Whitney test.

## 3. Results

### Distribution of CNS4 into plasma and brain compartments

To facilitate *in vivo* studies, we have carried out preliminary PK to determine the basic PK properties of CNS4 as presented in **Figure 1** and **Table 1**. CNS4 reaches maximum plasma and brain concentrations (T_max_) as quickly as 0.25 hours and was above the LOQ (T_min_) for up to 8 hours in the plasma and 12 hours in the brain tissue after a single intraperitoneal (IP) injection. Approximately 8.4% of the total (protein bound and unbound) plasma concentration is present in the brain tissue based on a comparison of the maximum concentrations (64.9 µg/mL vs 5.47 µg/ml) or 3.25% based on AUC_0-∞_ (52.6 hr*µg/mL vs 1.71 hr*µg/mL). Further, the volume of distribution (Vd/F) of 1.91 L/kg suggests that CNS4 is well distributed outside of the plasma compartment.

**Figure 1.**
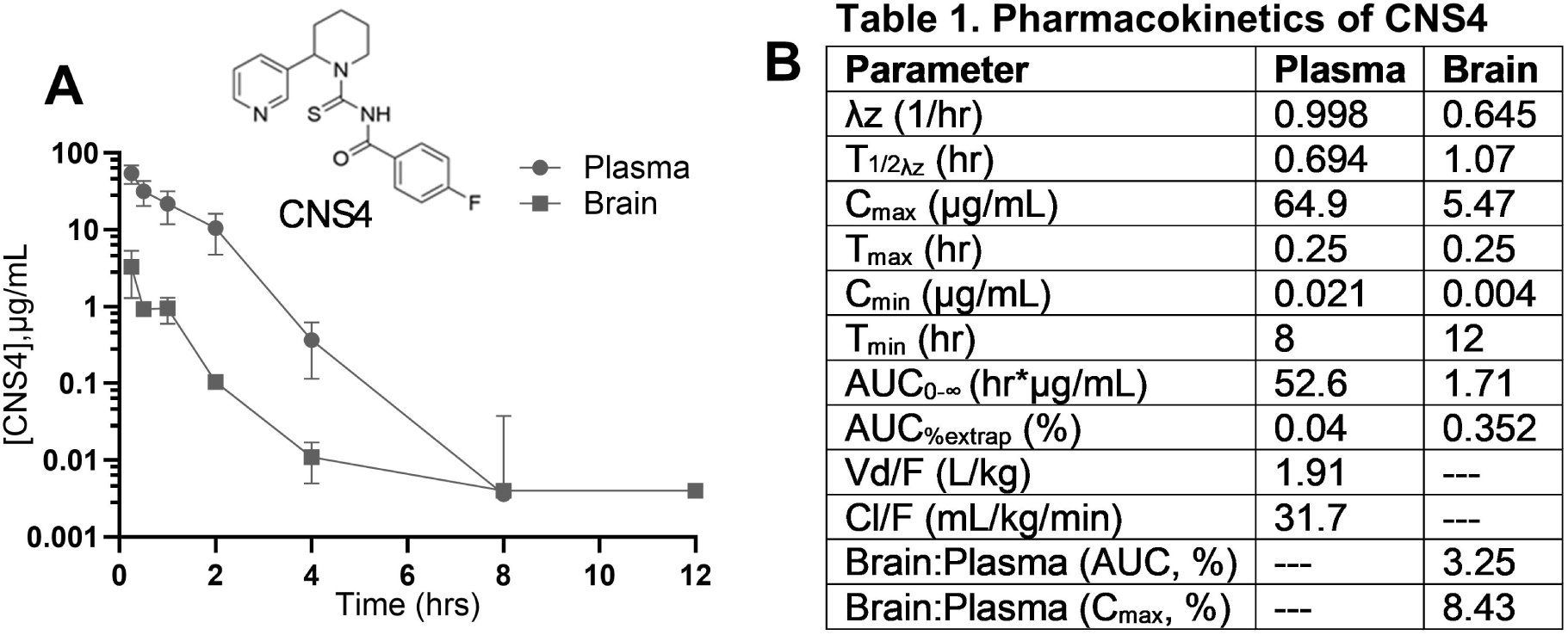
**CNS4 reaches plasma and brain tissue after intraperitoneal injection in male CF-1 mice**. Mice were euthanized, after 100mg/kg CNS4 dissolved in normal saline, in 0, 0.25, 0.5, 1, 2,4,8,12,24 and 48 hours. Limit of detection (LOD) was 0.25 ng analyte/mL plasma or brain tissue for all analytes. Plasma and brain sample analytes were within LOD until 8 and 4 hours respectively, as plotted (**A**). n=3 male CF-1 mice for each time point. Thirty mice for ten time points. Inset, chemical structure of CNS4. Data presented as mean ± sem. **B**. Table shows the calculated pharmacokinetic parameters: λ_z_ = Terminal rate constant. T ½ = Terminal-phase half-life. C_max_ = Maximum concentration. T_max_ = Time to maximum concentration. C_min_ = Minimum concentration. T_min_ = Time to minimum concentration. AUC_0–∞_ = Area under the concentration-time curve from time zero extrapolated to infinity. AUC_%extrap_ = Percentage of AUC that was extrapolated. VD/F = apparent volume of distribution for an extravascular route; Cl/F = apparent clearance for an extravascular route. Brain:plasma = percentage of drug measured in the brain versus the plasma based on AUC_0-∞_ or C_max_.

**Table 1.** Pharmacokinetics of CN54.

| **Parameter** | **Plasma** | **Brain** |
| --- | --- | --- |
| $\lambda_z$ (1/hr) | 0.998 | 0.645 |
| $T_{1/2\lambda_z}$ (hr) | 0.694 | 1.07 |
| $C_{max}$ ($\mu\text{g/mL}$) | 64.9 | 5.47 |
| $T_{max}$ (hr) | 0.25 | 0.25 |
| $C_{min}$ ($\mu\text{g/mL}$) | 0.021 | 0.004 |
| $T_{min}$ (hr) | 8 | 12 |
| $AUC_{0-\infty}$ (hr*$\mu\text{g/mL}$) | 52.6 | 1.71 |
| $AUC_{\%extrap}$ (%) | 0.04 | 0.352 |
| $Vd/F$ (L/kg) | 1.91 | --- |
| $Cl/F$ (mL/kg/min) | 31.7 | --- |
| Brain:Plasma (AUC, %) | --- | 3.25 |
| Brain:Plasma ($C_{max}$, %) | --- | 8.43 |

**Table 1. Pharmacokinetics of CNS4**
| Parameter | Plasma | Brain |
| --- | --- | --- |
| $\lambda_z$ (1/hr) | 0.998 | 0.645 |
| $T_{1/2\lambda_z}$ (hr) | 0.694 | 1.07 |
| $C_{max}$ (µg/mL) | 64.9 | 5.47 |
| $T_{max}$ (hr) | 0.25 | 0.25 |
| $C_{min}$ (µg/mL) | 0.021 | 0.004 |
| $T_{min}$ (hr) | 8 | 12 |
| $AUC_{0-\infty}$ (hr*µg/mL) | 52.6 | 1.71 |
| $AUC_{\%extrap}$ (%) | 0.04 | 0.352 |
| $V_d/F$ (L/kg) | 1.91 | --- |
| $Cl/F$ (mL/kg/min) | 31.7 | --- |
| Brain:Plasma (AUC, %) | --- | 3.25 |
| Brain:Plasma ($C_{max}$, %) | --- | 8.43 |

Mass spectrometry analysis of plasma and brain tissue samples identified ionization of molecules that match the molecular weight of metabolite-1 in both plasma and brain tissue and metabolite-2 only in plasma. Chemical structures of metabolites are provided in **Supplementary** Figure 1.

### Analgesic effect produced by CNS4 is different from an NSAID

With evidence that CNS4 reaches the expected site of action, we have started preclinical studies. To investigate the *in vivo* analgesic effect of CNS4, we first performed a hot plate analgesic assay. Results from this experiment revealed that CNS4 increased mice escape latency (paw-licking) by 1.74-fold (12.01 ± 1.81 vs 6.90 ± 0.46s, p=0.04, Kruskal-Wallis and post hoc Dunn’s multiple comparisons test) compared to the saline control. Whereas, meloxicam exhibited 1.09-fold (7.56 ± 0.75 vs 6.90 ± 0.46, p=0.99) compared to saline (**Figure 2**). A 55°C hotplate was used to increase the central nervous system involvement. Meloxicam is a non-steroidal anti-inflammatory drug that has no central analgesic effect (Engelhardt, 1996). Thus, meloxicam’s effect was insignificant. This data also supports the finding from PK assay that CNS4 reaches brain tissue, and CNS4’s effect is different from that of peripherally acting non-steroidal anti-inflammatory drug meloxicam. To assess the similarity between mice obtained from two different providers, we compared the hot plate assay escape latency of the saline-treated (control) groups from Phase 1 (CRL) and Phase 2 (JAX). The results showed no statistically significant difference in escape latency between the two groups (CRL: 7.66 ± 0.66 s vs. JAX: 6.14 ± 0.43 s; n = 7; p = 0.21; unpaired, non-parametric Kolmogorov-Smirnov test).

**Figure 2.**
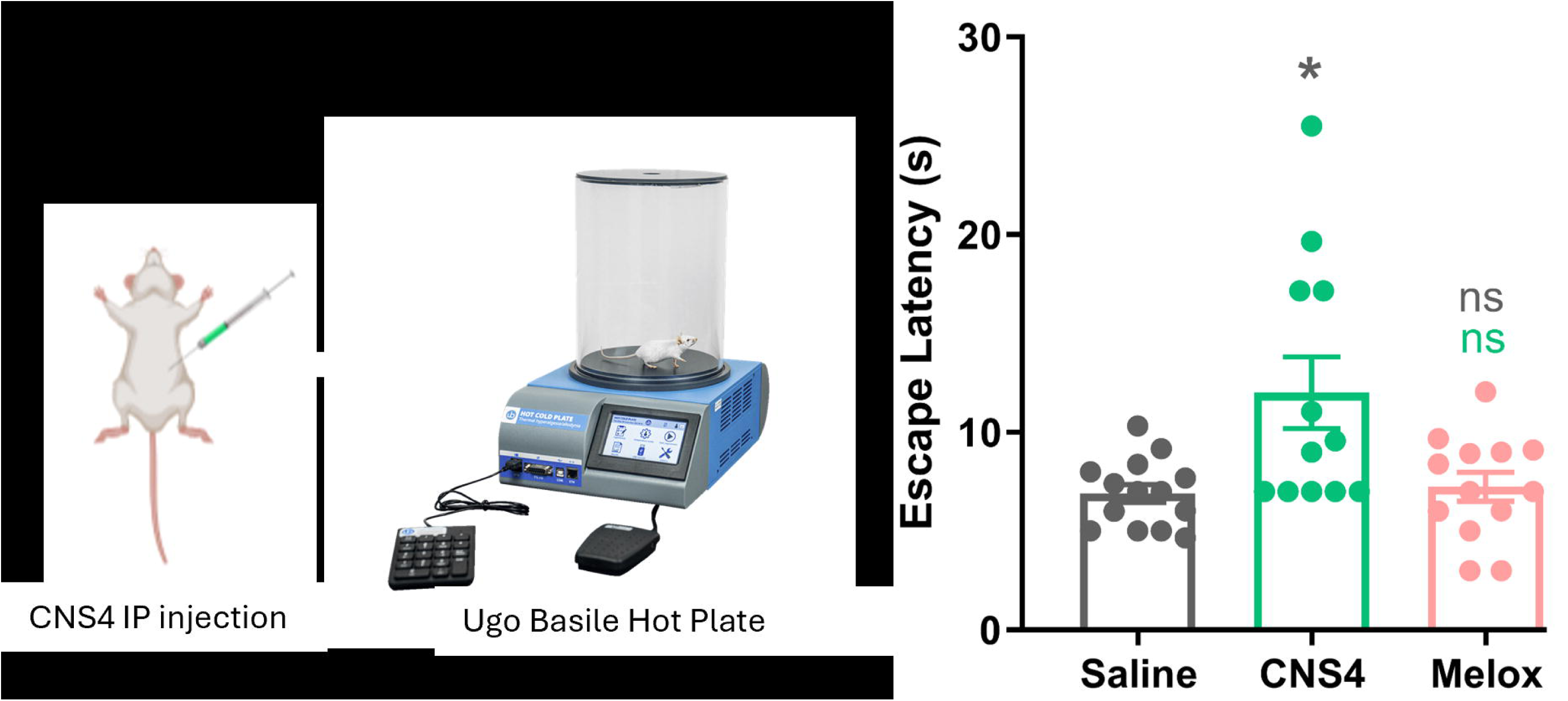
Delayed paw-licking response in CNS4-treated mice during the hot plate test. Histograms show paw licking (escape) latency for saline (6.90±0.46s), CNS4 (12.01±1.81s), and meloxicam (7.56±0.75s) in 55°C hotplate after IP administration of saline or 100mg/kg CNS4 or meloxicam (melox) 5mg/kg. Dots represent sample size [saline 14(8m,f6), CNS4 12(6m,6f), melox 14(6m,8f)] of C57BL male (m) or female (f) mice. Groups are compared by Kruskal-Wallis test and post hoc Dunn’s multiple comparisons test. *p=0.043. Data presented as mean ± sem. ns, not significant, p>0.05. Asterisk or ns color represents the comparison group.

### Effect of CNS4 on foot shock sensation and fear memory

Since CNS4 increased thermal escape latency, we hypothesized that it might also reduce the sensitivity to foot shock and, therefore, influence fear memory formation in the contextual fear conditioning experiment. A flow chart diagram (**Figure 3A&B**) shows the FC training and testing protocol. FC assay results revealed that on paired CS and US (day 1) experiments, CNS4-treated animals (both males and females) experienced anecdotally less shock compared to the saline-treated group [**Supplementary** Figure 2 (video)].

**Figure 3.**
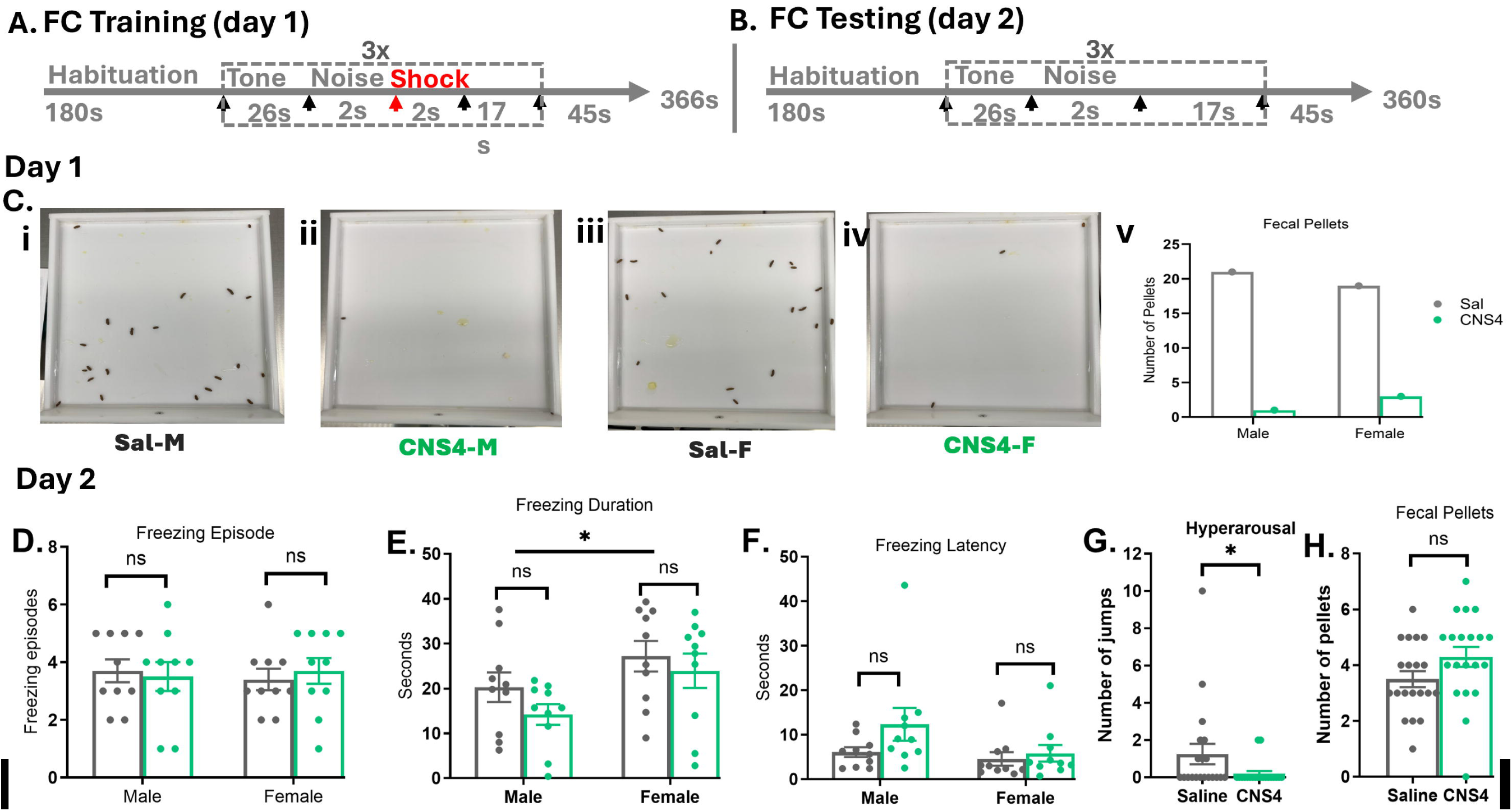
Effect of CNS4 on electric shock-induced fear conditioning (FC) on male and female mice. Flow chart for fear condition (FC) training (**A**) and testing (**B**) experiment. Tone, volume=50,duration=26s, frequency=4kHz; White noise, volume=50, duration=2s, 20kHz; Shock, 0.5mA, 2s. On testing (day 2), mice underwent the same protocol but no shock. **Ci–iv** displays the fecal pellet images captured after each group’s FC training on day 1, with the histograms (v) representing a numerical comparison of the results. . Freezing episode (**D**), duration (**E**), latency (**F**), hyperarousal (**G**), and the number of individual mouse fecal pellets data (**H**) collected on FC testing day, are compared between saline and CNS4 treated mice. Freezing data sets were initially analyzed by Two-Way ANOVA, then Uncorrected Fisher’s Least Significance Difference test, to find out differences between individual groups. Post hoc tests are marked as downward pointing edges and the Two-way ANOVA comparison is marked as a straight line on top of the bars. Data points represent the biological replicates, n=10 mice per group and four groups (Sal-M, Sal-F, CNS4-M, CNS4-F). Hyperarousal (**H**) Fecal pellets (**G**) are analyzed by Mann Whitney test between male and female mice, n=10 per group. Data presented as mean ± sem. ns= not significant, p>0.05; *=p<0.05; **=p<0.01; ***=p<0.001. **D, E &F** show the freezing data from 45 seconds (181-225s) after hearing the first tone on FC testing day. The second (227-271s) and third (272-316s) tone freezing data is provided as supplementary Figure 3.

Analysis of fecal pellet data obtained after day 1 stimulus protocol revealed that CNS4-treated male (1 vs 21) and female (3 vs 19) mice defecated numerically less compared to the saline-treated groups (**Figure 3Ci-v**). This data suggests that CNS4-treated mice probably did not feel the shock as much as saline-treated mice. We thought CNS4 could have produced an anesthetic effect, as drugs acting on NMDAR are dissociative general anesthetics. If CNS4 produced cognitive dysfunction, on day 2, the CNS4-treated mice might not remember the CS and might not have paired it with the US. In contrast to expectation, analysis of freezing data obtained after tone 1 (181-225s), tone 2 (227-271s) and tone 3 (272-316s) of the FC testing day revealed that CNS4 had no impact on the freezing parameters. The number of freezing episodes (m, 3.70 ± 0.40 vs 3.50 ± 0.50, p=0.77; f, 3.40 ± 0.37 vs 3.70 ± 0.45, p=0.66), freezing duration (m, 20.27 ± 3.31s vs 14.22 ± 2.30s, p=0.56; f, 27.20 ± 3.40s vs 23.93 ± 3.84s, p=0.89), and freezing latency (m, 6.07 ± 1.09s vs 12.34 ± 3.68s, p=0.22; f, 4.56 ± 1.51s vs 5.83 ± 1.87s, p= 0.97) after tone 1 indicate that the freezing parameters of CNS4-treated mice were indifferent from the saline-treated mice (**Figure 3D-F**), as determined by two-way ANOVA and post hoc Uncorrected Fisher’s Least Significance Difference test. Notably, a comparison of data across the sexes (row factor) using two-way ANOVA indicates that the freezing duration of female mice is longer than the male mice (25.5 ± 2.52s vs 17.25 ± 2.07s, p=0.015). Tone 2 & 3 data are similar to tone 1 and they are provided in **Supplementary** Figure 3.

### Phenotypes associated with fear improved in CNS4-treated mice

Since the data obtained from the FC suggested no change in fear memory, we hypothesized that CNS4 might not have altered the behavioral phenotypes associated with FC. In contrast, the hyperarousal data reveals that CNS4-treated mice made fewer jumps compared to the saline-treated ones (0.20 ± 0.13 vs 1.25 ± 0.54, p=0.04, Mann Whitney test) on FC testing day when there was no shock, **Figure 3G**. Next, we studied the number of fecal pellets for each animal after their time in the FC chamber on testing day. Analysis of the number of fecal pellets indicates that CNS4-treated mice defecated a comparable number of fecal pellets as saline-treated mice (4.30 ± 0.35 vs 3.50 ± 0.29 pellets, p=0.054, Mann Whitney test), **Figure 3H**. No difference was identified when comparing all four groups (Sal-M, CNS4-M, Sal-F and CNS4-F) based on sex (p=0.99) or treatment (p=0.09) using the two-way ANOVA test. All animal fecal pellet pictures are provided as **Supplementary** Figure 4.

### CNS4 did not alter locomotion in male or female mice

To compare the individual animals’ fear response with their baseline activity, an open field test was performed before (baseline) and after the FC experiment. The only difference (Δ) in locomotion that was identified between males and females (row factor) is on the distance traveled (m,6.42 ± 2.14 vs f, 12.01±1.36 meters, p=0.02, Two-way ANOVA), **Figure 4A**. However, this difference was insignificant when compared with sex-matched treatments (p>0.05, post hoc Šídák’s multiple comparisons test). On the other hand, there was a significant difference between saline and CNS4 treated groups (column factor) in distance traveled (saline, 6.81 ± 1.82 vs CNS4, 11.8 ± 1.87 meters, p=0.04), mobile episodes (saline, −3.67 ± 1.09 vs CNS4, −6.57 ± 0.75, p=0.03), and time immobile (saline, −14.98 ± 5.02 vs CNS4, −36.93 ± 4.66, p=0.0027). However, none of the sex-matched treatment group comparisons, made by post hoc Šídák’s multiple comparison tests, were significant (p>0.05).

**Figure 4.**
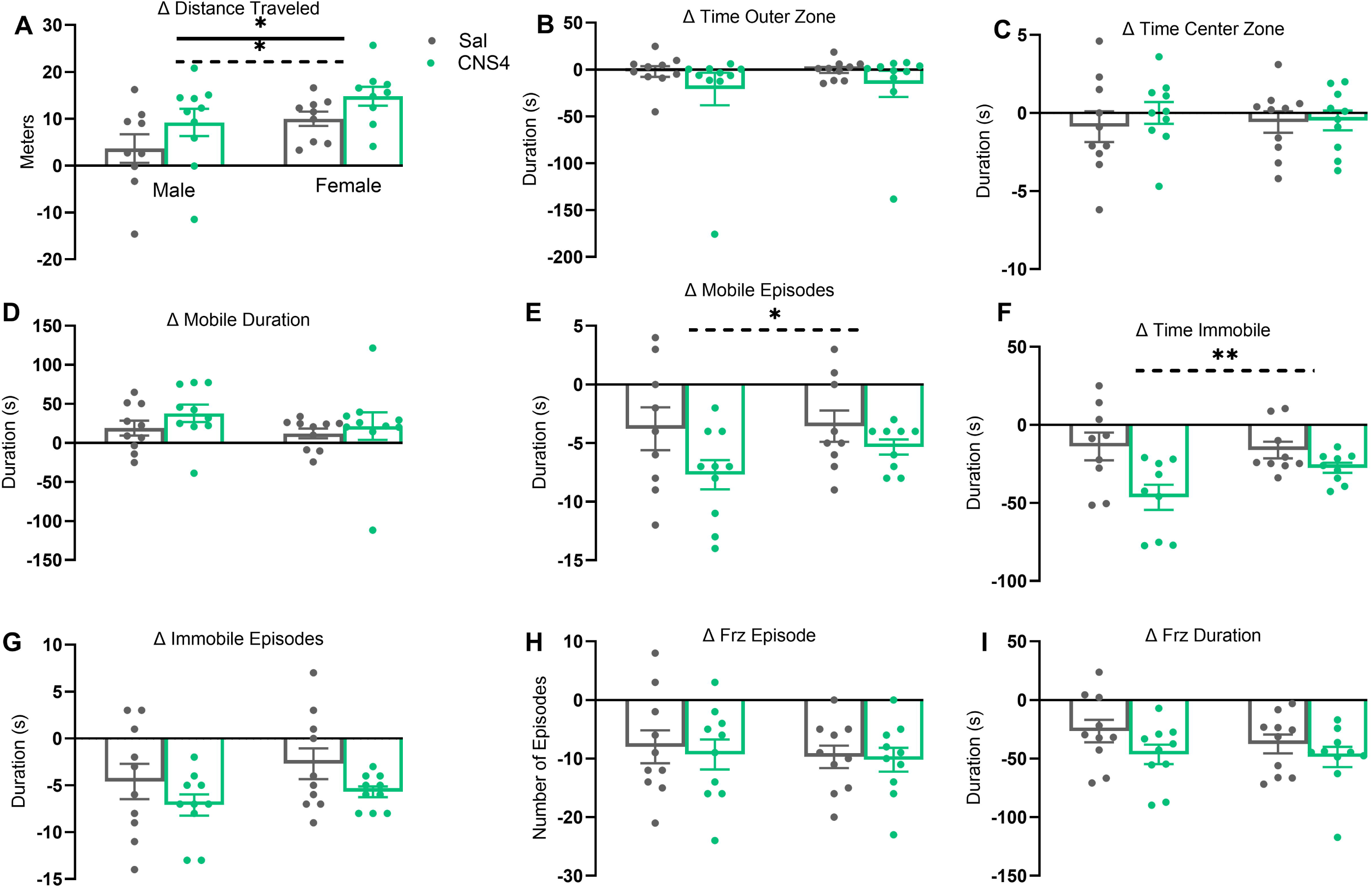
Effect of CNS4 on locomotion and anxiety before and after the FC experiment. Histograms (**A–I**) indicate the difference (Δ) between the baseline and after-treatment data for each animal on various locomotion and anxiety-related behavior parameters (as labeled) studied from the open field assay. Mice did not receive saline or CNS4 before the FC experiment; they did receive saline (Sal) or CNS4 fifteen minutes before the FC training day (day 1). There were four groups of animals [control male (Sal-M), CNS4 treated male (CNS4-M), control female (Sal-F) and CNS4 treated female (CNS4-F)] and each group had ten (n=10) mice. All parameters were initially analyzed by Two-Way ANOVA, then post hoc Šídák’s multiple comparisons test, to find out differences between individual groups. Significant differences between males and females (row factor –marked with line), and saline and CNS4 treated groups (column factor –marked with dotted line) are marked on the histograms. No parameter was found to be significantly different between sex-matching saline and CNS4 treatment by the post hoc Šídák’s multiple comparisons test. Data points represent biological replicates. Data presented as mean ± sem. The data that are not significant (p>0.05) are not labeled. *=p<0.05; **=p<0.01.

### CNS4 prevents excess sucrose intake in male and female mice

Further, we studied CNS4-induced changes in sucrose preference in male and female mice. Baseline SPT revealed a three to four-fold increase in 1% w/v sucrose in water solution consumption compared to plain water in all four cages (Sal-M, 4.2x; CNS4-M, 4.8x; Sal-F, 4.1; CNS4-F, 3.4x), **Figure 5A**. Consistently, two days after FC training, all four groups again preferred sucrose (Sal-M, 5.9x; Sal-F, 9.0x; CNS4-M, 2.9; CNS4-F, 3.3x), **Figure 5B**. However, comparing fold changes before and after FC revealed that Sal-M and Sal-F groups consumed 1.4x and 2.9x more sucrose after FC, while CNS4-M and CNS4-F groups consumed a little less (0.6x) and about the same (0.97x) amount of sucrose after FC, respectively, **Figure 5C**.

**Figure 5.**
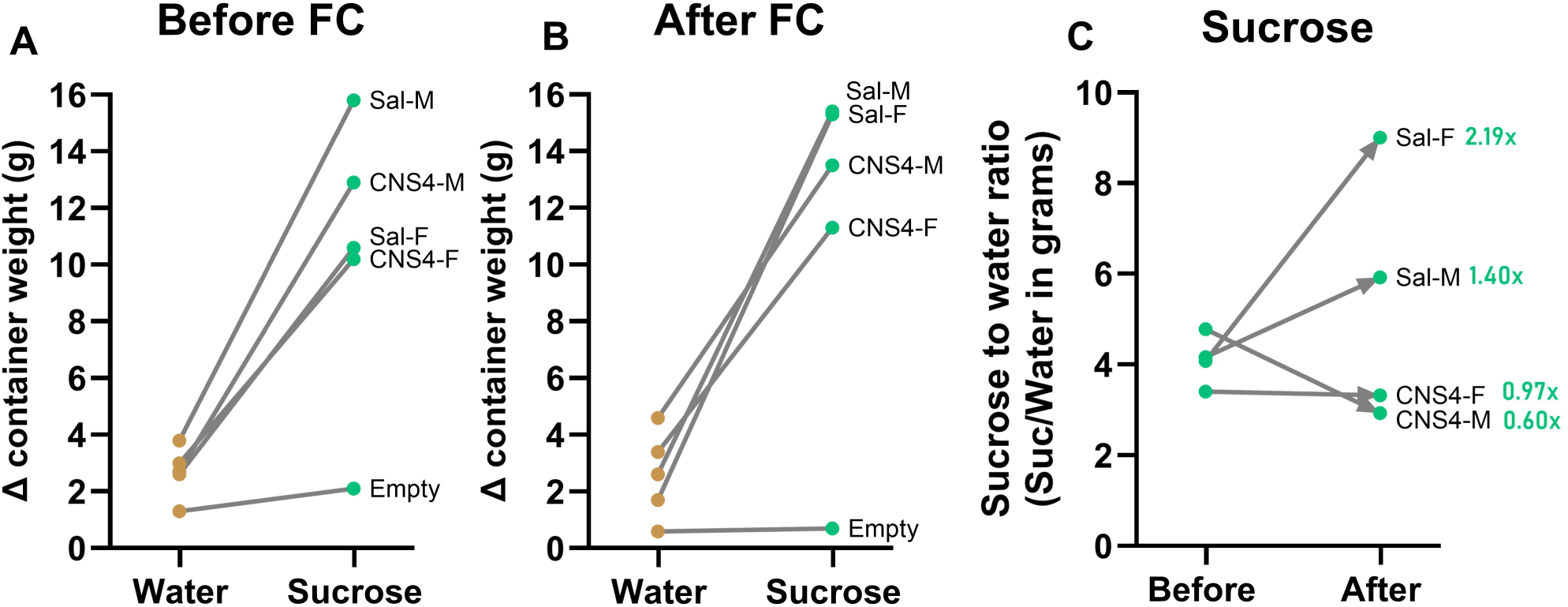
Effect of CNS4 on sucrose preference in mice after FC experiment. Pre-weighed water and 1% w/v sucrose in tubes were provided on each cage of four groups of mice before (**A**) and after (**B**) the fear conditioning (FC) experiment. In **A&B**, the y-axis represents the difference (Δ) in container weight. **C**, Comparison of before and after FC sucrose to water ratio. Data points were obtained by dividing the sucrose by water values presented before (A) and after (B) FC. Numbers by the labels indicate the fold difference. Empty shows the data obtained from the cages where no animal was present.

## Discussion

Based on the predicted physicochemical properties and unique pharmacodynamic characteristics of CNS4 (Boehringer et al., 2023; Costa, 2018; Costa et al., 2021), we have carried out a translational pharmacology study. The preliminary PK work revealed that CNS4 reaches maximum concentration in plasma and brain tissue as rapidly as fifteen minutes after a single intraperitoneal injection (**Table 1**). The rapid T_max_ in the brain tissue suggests that CNS4 distributes simultaneously to both compartments following a concentration gradient. The highly lipophilic nature of CNS4 (LogP _o/w_ =2.42) supports rapid passive diffusion across the BBB; however, an active transport mechanism may also be present. This study reports total concentrations and not unbound (active) drug concentrations, which requires further study, as does further quantification of metabolites. However, these initial findings confirm distribution of the drug from the plasma into the CNS, the presumed site of action. These results motivated us to carry out a subsequent hot plate assay to study the analgesic effect of CNS4.

Results from the hot plate assay revealed that CNS4 increases escape latency, as an indicator of analgesic effect. Further, this effect differs from that of a peripherally acting non-steroidal anti-inflammatory drug, meloxicam. Collectively, data obtained from the PK and hotplate assays suggest that CNS4 is crossing the BBB, reaching brain tissue, and producing a central analgesic effect. While this data does not necessarily confirm that the observed effect is an outcome of CNS4 directly binding with the neuronal NMDARs, it is known from previous *in vitro* experiments that CNS4 can directly potentiate or inhibit NMDAR subtypes based on agonist concentration (Boehringer et al., 2023; Costa et al., 2021). Since there is a potential for target engagement, we then studied the effect of CNS4 on contextual FC as there is a compelling but enigmatic role of NMDAR subtypes in various components of post-traumatic stress disorders (PTSD) (Dalton et al., 2012; Dubois et al., 2016; Dubois and Liu, 2021; Gao et al., 2010; Leaderbrand et al., 2014; Rasmussen, 2015; Salimando et al., 2020). Moreover, both PTSD and chronic pain involve alterations in brain regions related to emotion, cognition, and sensory processing (Burris et al., 2009; Jenewein et al., 2009; Sharp and Harvey, 2001). Chronic pain complaints are common in patients with a primary diagnosis of PTSD, with prevalence rates estimated to be as high as 80% (Lew et al., 2009; Otis et al., 2009).

Psychological and experimental evidence suggest that defecation is an indication of heightened fear and anxiety (Flint et al., 1995; Hall, 1934; Ramos, 2008; Russo and Parsons, 2021). This theory reveals that CNS4-treated mice did not experience fear and anxiety on FC training (day1) when compared to the saline-treated mice (fecal pellets: m, 1 vs 21; f, 3 vs 19). This drastic difference can be an indication of potent analgesic or general anesthetic activity of CNS4. Indeed, a non-competitive NMDA receptor antagonist, ketamine, is a clinically used dissociative general anesthetic agent (Domino et al., 1965). Further, many opioid and non-opioid drugs like ketamine produce analgesic effects at lower doses and sedation at higher doses (Bell and Kalso, 2018; Cohen et al., 2024; Guldner et al., 2006). We have used a 100mg/kg dose, which can be interpreted as a relatively high dose. Therefore, we intended to prove that mice had full consciousness inside the chamber, and they associated the CS with the US. Although it can be visually observed from the video file [**Supplementary** Figure 2 **(video)**] that the mice were not asleep, but rather were moving in the chamber and experiencing shock to some extent, this does not rule out the possibility of dissociation from the environment. However, the contextual fear memory results provided a crucial input to circumvent this assumption . Fear memory formation requires animals to remember the context in which they receive adverse stimuli, and this is dependent on the hippocampus, where GluN1/2AB triheteromeric NMDAR are predominately expressed (Biedenkapp and Rudy, 2007; Davis, 1986; Kim and Jung, 2006; Lafon-Cazal et al., 2002; Phillips and LeDoux, 1992; Soares and Lee, 2013; Stroebel et al., 2018; Tovar et al., 2013). Results from FC testing (day 2) reveal that there was no change in the freezing episode, duration, or latency in CNS4-treated mice compared to the saline-treated male or female mice after any of the three tones studied, indicating that there was no significant difference in the hippocampus-dependent contextual memory needed to associate CS with US.

However, the number of jumps observed on FC testing day, when there was no shock, suggests that CNS4 significantly reduced hyperarousal (a total of 25 vs 4 jumps). Hyperarousal is an indicator of heightened fear and anxiety (Cohen and Zohar, 2004; LeDoux, 2000). Previous studies demonstrated that blockade of NMDAR in the basolateral amygdala prevents fear extinction since NMDAR-mediated long-term potentiation in the basolateral amygdala and thalamus is critical for extinction learning (Blair et al., 2001; Johansen et al., 2011; Maren, 1999). Further, a growing body of evidence suggests that impairment in acquiring fear extinction is a key mechanism of PTSD, and mice lacking the GluN2D subunit of NMDAR were unable to extinguish fear memory (Dubois and Liu, 2021; Ogden et al., 2014; Salimando et al., 2020). In contrast, low-dose NMDAR blockers reduced hyperarousal and depressive symptoms of PTSD in humans (Albott et al., 2018; Battista et al., 2007; Feder et al., 2014). Therefore, we speculate that CNS4-induced partial negative allosteric modulation of GluN1/2D and minimal positive allosteric modulation of 1/2AB triheteromeric NMDAR (Costa et al., 2021), particularly when activated by high concentrations of glutamate in the amygdala and thalamus, which occurs with heightened fear (Simic et al., 2021; Walker and Davis, 2002), might play role in reducing hyperarousal. Moreover, this could be a fear extinction induction component.

The comparable number of fecal pellets in CNS4-treated mice on FC testing day (4.30 ± 0.35 vs 3.50 ± 0.29 pellets, p=0.054) indicates normal food intake and overall normal fluid intake . Remarkably, this data is opposite to what was observed on FC training day (day1) when CNS4-treated mice defecated a lot fewer pellets. While connecting the number of fecal pellets directly with food intake or fear, we also recognize that CNS4 might have altered gastrointestinal motility. This needs to be addressed in future studies.

Stress and reward-seeking behavior have a complex relationship based on multiple contributing factors (Mather and Lighthall, 2012). Uncontrollable stress leads to anhedonic behavior and dysfunction within mesocorticolimbic dopaminergic pathways implicated in incentive motivation and hedonic coding (Anisman and Matheson, 2005; Henn and Vollmayr, 2005). In contrast, acute stress influences long-term reward maximization (Byrne et al., 2019; Mather and Lighthall, 2012). Comparison of sucrose preference before and after FC suggests an increase in sucrose preference after the FC training in saline-treated mice (male 1.4x and female 2.2x), indicating either reward-seeking behavior or stress-mediated increased metabolic process needing more sucrose, or both. CNS4-treated mice exhibited minimal or largely unchanged (male 0.6x and female 0.97x) sucrose preference after FC. Possibly, the less aversive experience of foot shock in CNS4-treated mice prevented the consequential development of reward-seeking behavior that was observed in the saline-treated mice about fifty hours after the FC training. Further, this data reinforces the potent analgesic effect of CNS4.

Aberrant changes in synaptic glutamate concentrations are implicated in the etiology of clinical symptoms of depression and many other neuropsychiatric conditions (McGrath et al., 2022; Moussawi et al., 2011; O’Donovan et al., 2017). The GluN2D subtype of NMDAR plays a critical role in the pathogenesis of these conditions due to its peculiar characteristics, including slow

deactivation, extrasynaptic localization, expression in both excitatory and inhibitory neurons, and developmental regulation (Hansen et al., 2017; Paoletti et al., 2013; Wyllie et al., 2013). Therefore, drugs that modulate GluN2D are of major clinical and pharmaceutical interest (Paoletti et al., 2013). In the past fifteen years, numerous compounds have been identified to selectively modify the function of GluN2D receptors with the goal of translating them into clinically useful drugs (Callahan et al., 2020; Costa et al., 2009; Costa et al., 2010; Mullasseril et al., 2010; Perszyk et al., 2020; Yi et al., 2020). However, to the best of our knowledge, none of the compounds that are primarily acting on GluN2D reached human clinical trials. One reason is the lack of translational efforts. In the present study, we have successfully translated an *in vitro*-level NMDAR modulator to rodents and proved its pharmacological effects, which are potentially associated with the modulation of native NMDA receptors. Future preclinical investigations of CNS4 and its analogs will help develop clinically useful drugs for pain associated with stressful conditions.

## Acknowledgments

Authors thank Virginia Tech animal house staff for taking care of our animals, and Jessica Muller for reading the manuscript.

## Data availability statement

All data that supports the findings of this work is provided in the manuscript and in the supplementary documents. Raw data can be obtained from the corresponding author with a reasonable request.

## Authorship Contribution

Conceptualization and participated in research design: Costa

Conducted experiments: Costa, Hines, Phillip, Boehringer, Council-Troche

Performed data analysis: Costa, Anandakrishnan and Davis

Wrote or contributed to writing the manuscript: Costa and Davis

## Footnotes

This work was supported by the VCOM Research Eureka Accelerator Program (REAP) Grant funded to BMC.

