## Supplementary Figure 1 for "Preliminary pharmacokinetics and *in vivo* studies indicate analgesic and stress mitigation effects of a novel NMDA receptor modulator"

**Supplemental Figure 1**

Article title: Preliminary pharmacokinetics and in vivo studies indicate analgesic and stress mitigation effects of a novel NMDA receptor modulator

Author list: Blaise M. Costa, De'Yana Hines, Nakia Phillip, Seth C. Boehringer, Ramu Anandakrishnan, McAlister Council-Troche, Jennifer L. Davis

Journal title: The Journal of Pharmacology and Experimental Therapeutics

Manuscript number: JPET-D-24-00077R1

**Predicted Metabolites of CNS4**

**Metabolite-1**

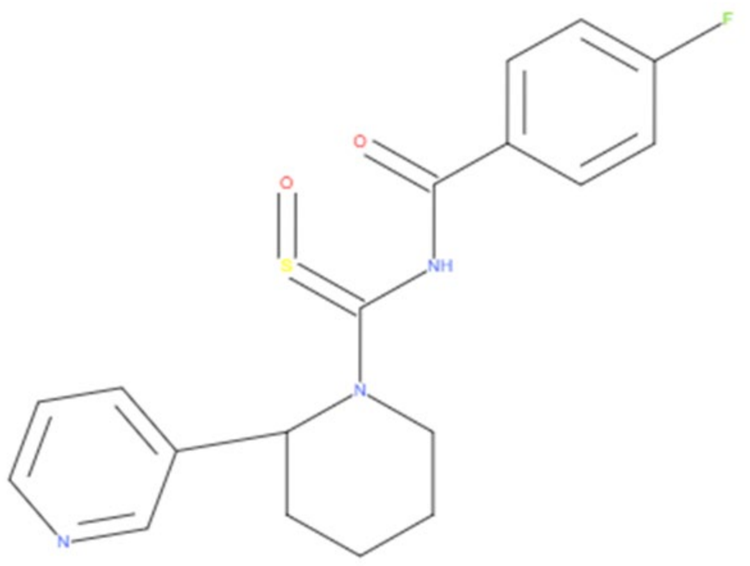


Rank 1 (Score: 0.41)

**Molecular weight 359.42 g/mol**

**Smile: O=S=C(N1CCCCC1c1cccnc1)NC(=O)c1ccc(cc1)F**

**Reaction: Oxidation (Phase 1)**

**Supplemental Figure 1**

Metabolite-2

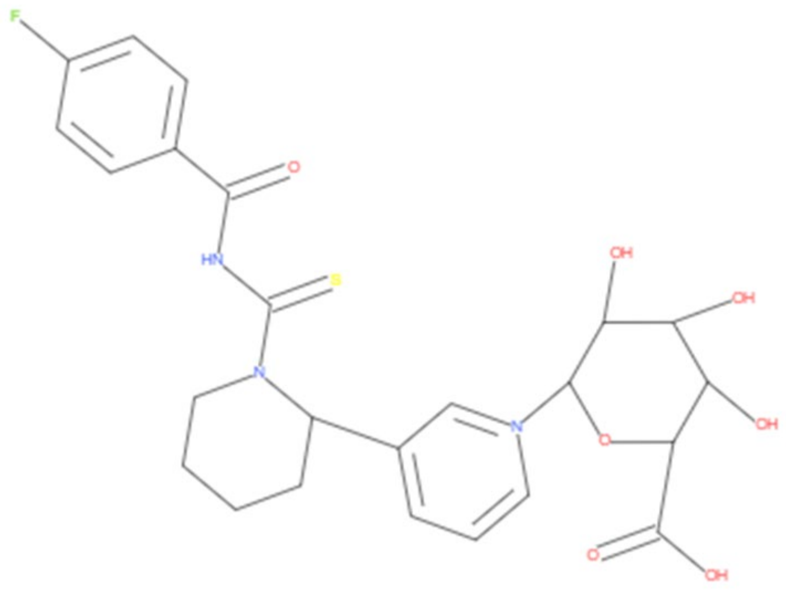

Rank 2 (Score: 0.38)

**Molecular weight 520.55 g/mol**

**Smile: Fc1ccc(cc1)C(=O)NC(=S)N1CCCCC1c1ccc[n+](c1)C1OC(C(=O)O)C(C(C1O)O)O**

**Reaction: Glucuronidation (Phase 2)**
