## Supplementary figures and images for "Preliminary pharmacokinetics and *in vivo* studies indicate analgesic and stress mitigation effects of a novel NMDA receptor modulator"

### Supplementary Figure 3

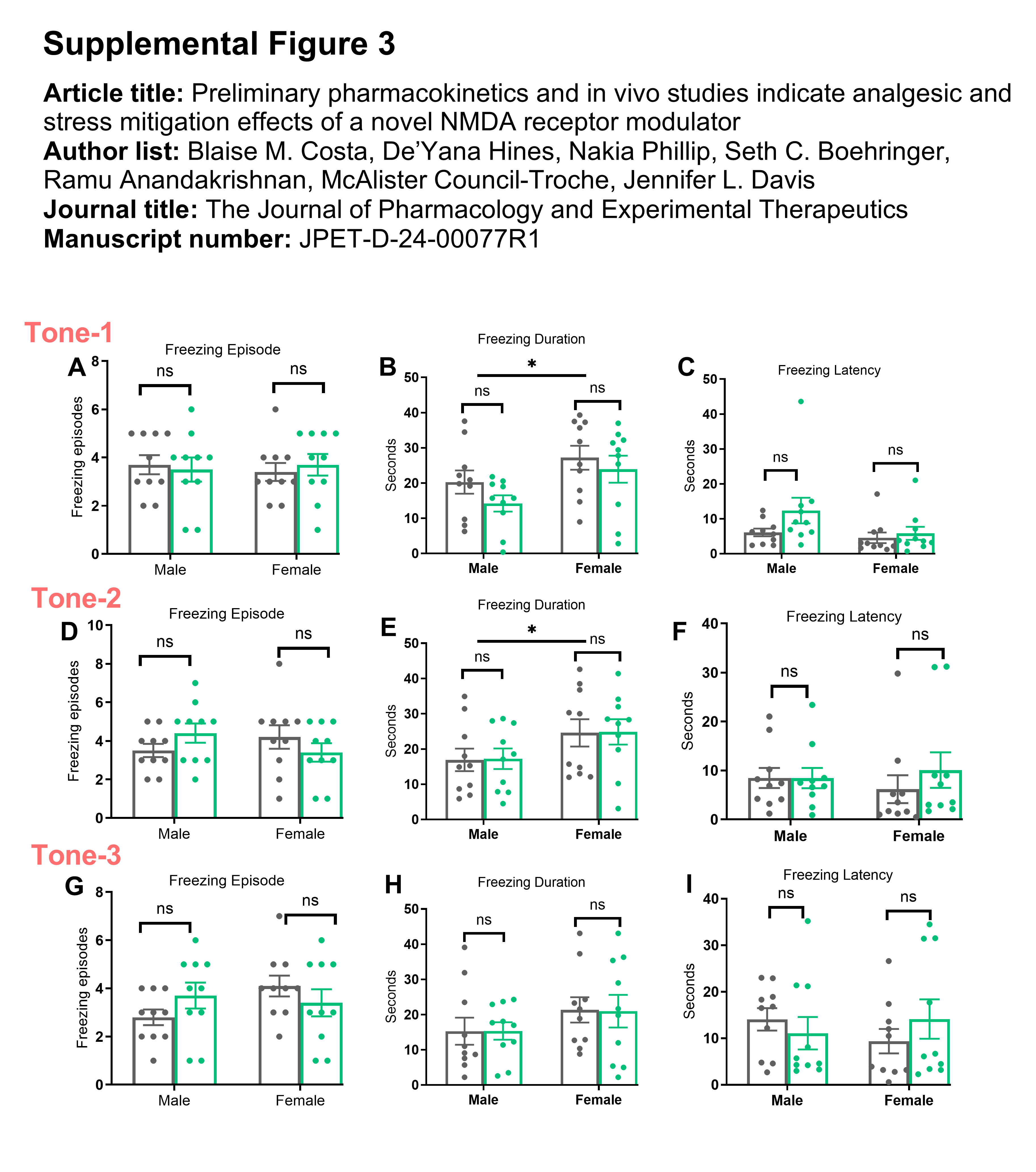
