## Supplementary Figure 4A for "Preliminary pharmacokinetics and *in vivo* studies indicate analgesic and stress mitigation effects of a novel NMDA receptor modulator"

Pictures are arranged in the following order: Control male 1-5, control female 1-5, CNS4 male 1-5, and CNS4 female 1-5.

Article title: Preliminary pharmacokinetics and in vivo studies indicate analgesic and stress mitigation effects of a novel NMDA receptor modulator

Author list: Blaise M. Costa, De'Yana Hines, Nakia Phillip, Seth C. Boehringer, Ramu Anandakrishnan, McAlister Council-Troche, Jennifer L. Davis

Journal title: The Journal of Pharmacology and Experimental Therapeutics

Manuscript number: JPET-D-24-00077R1

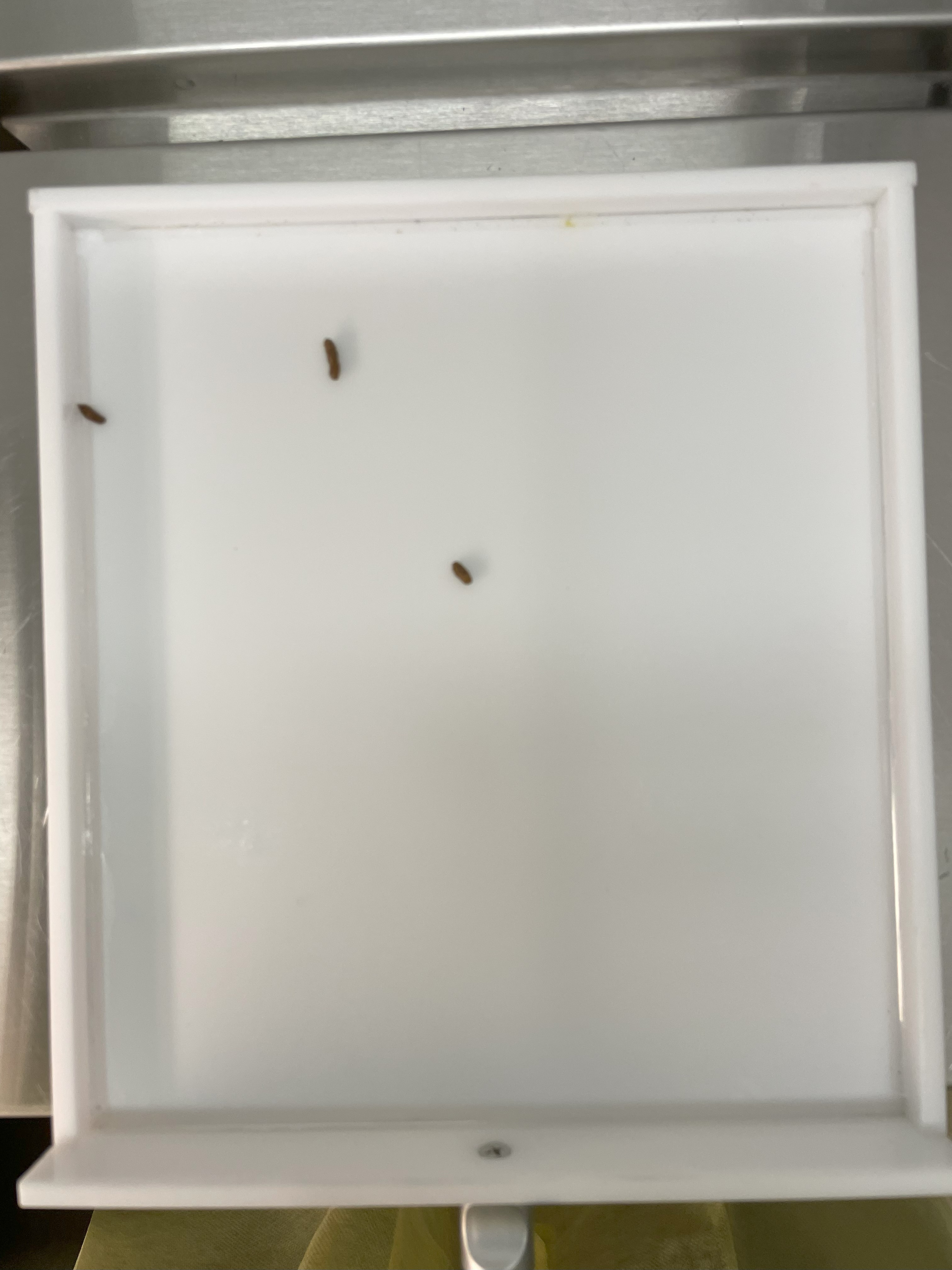

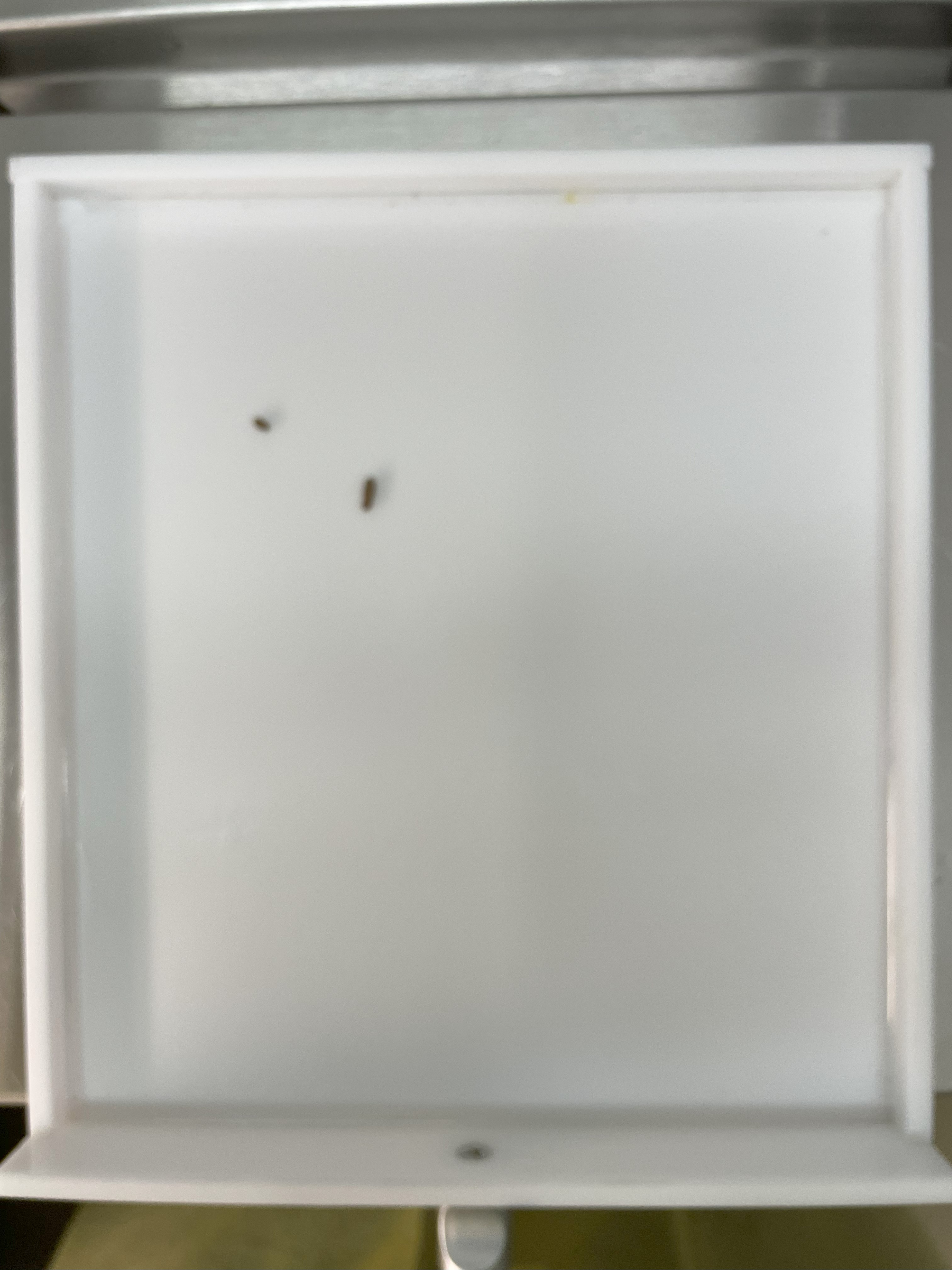

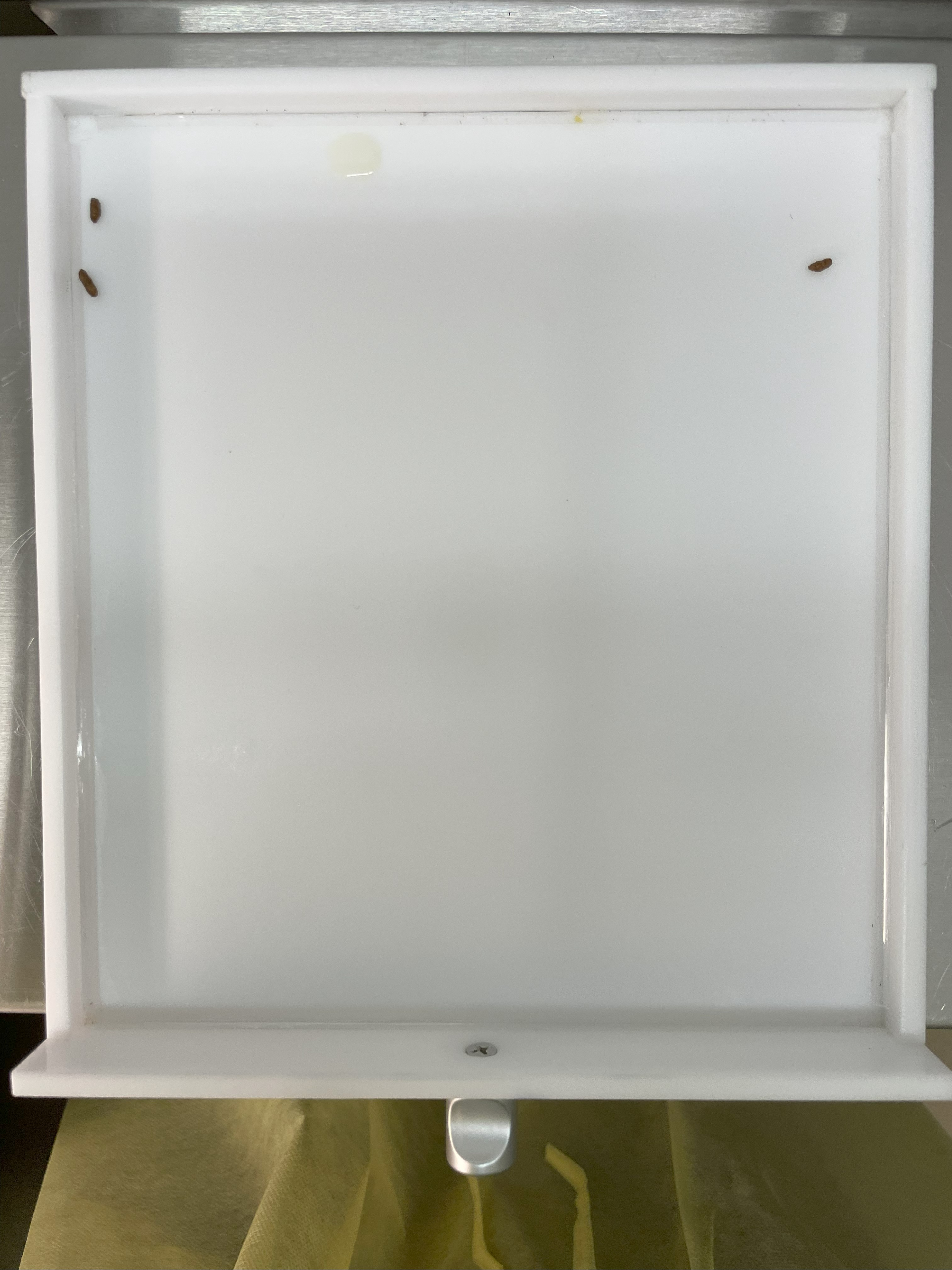

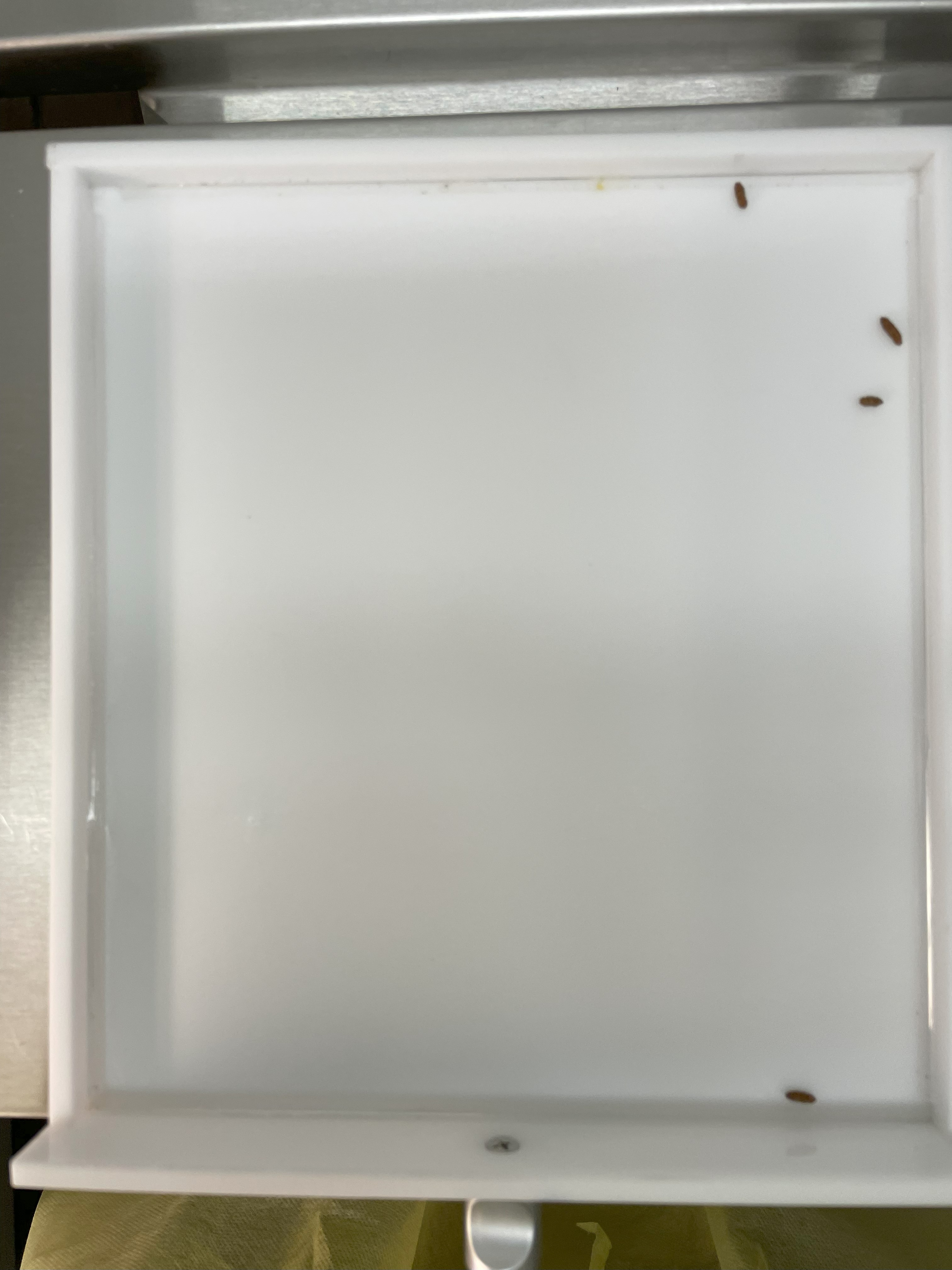

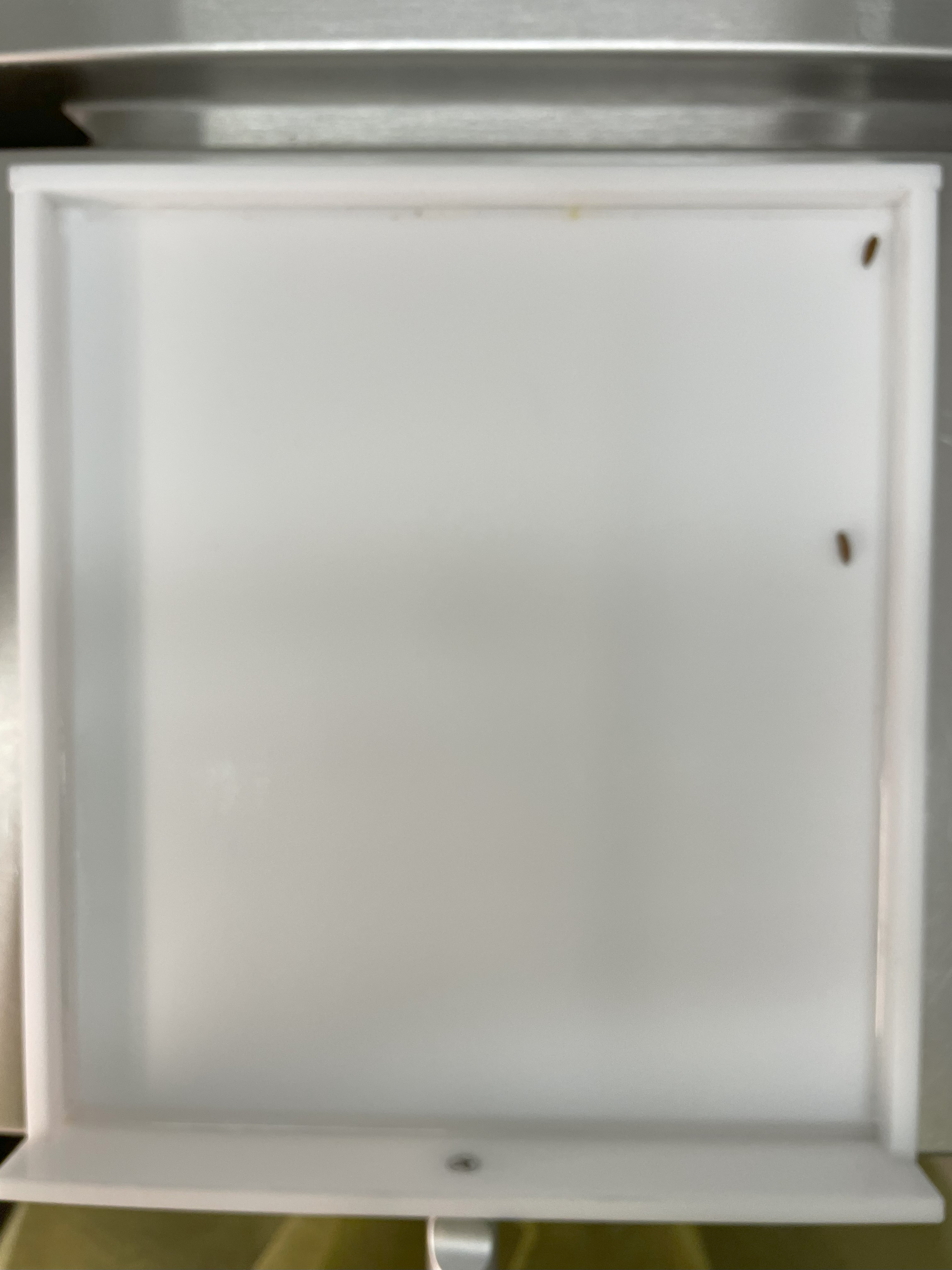

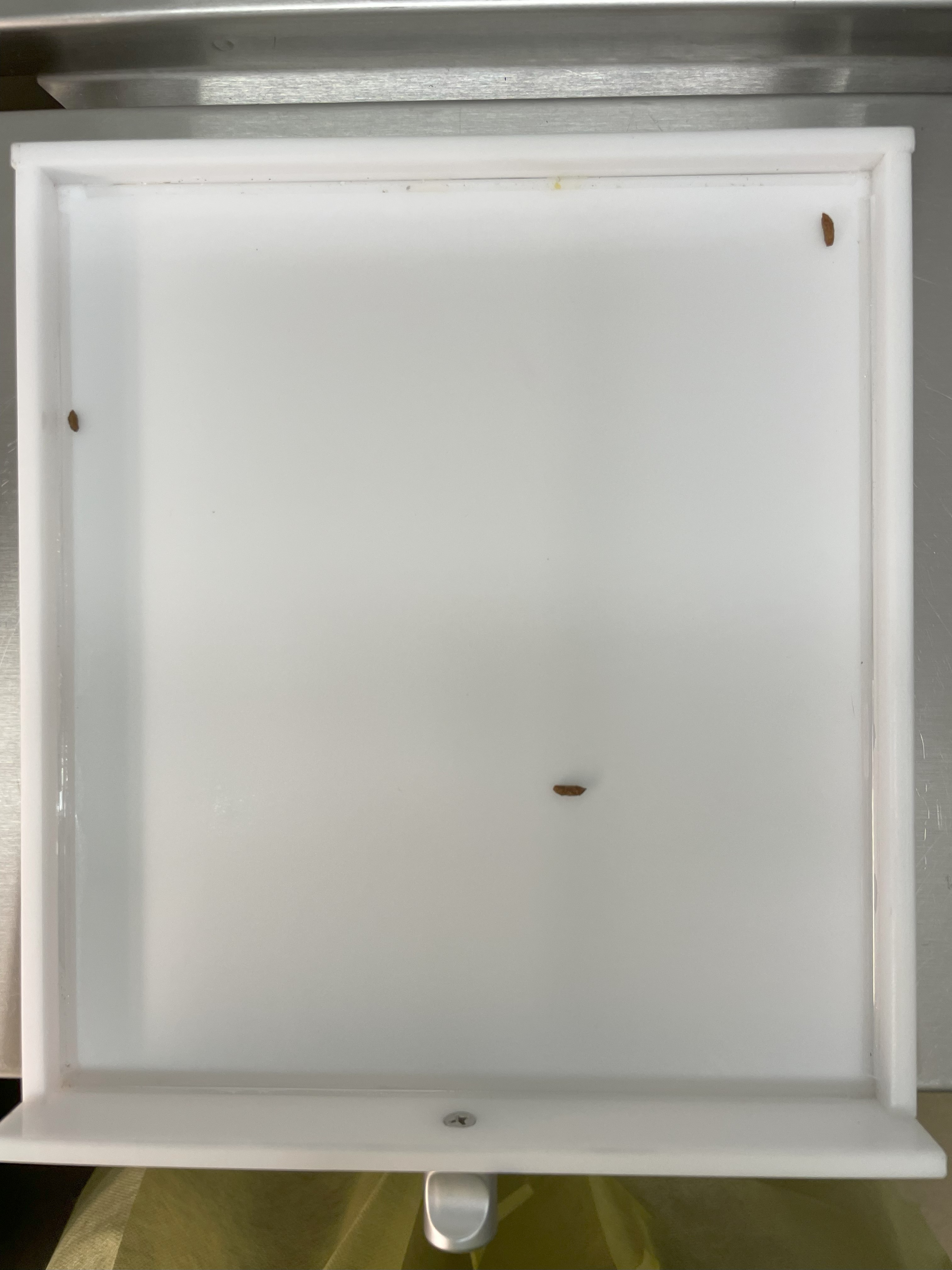

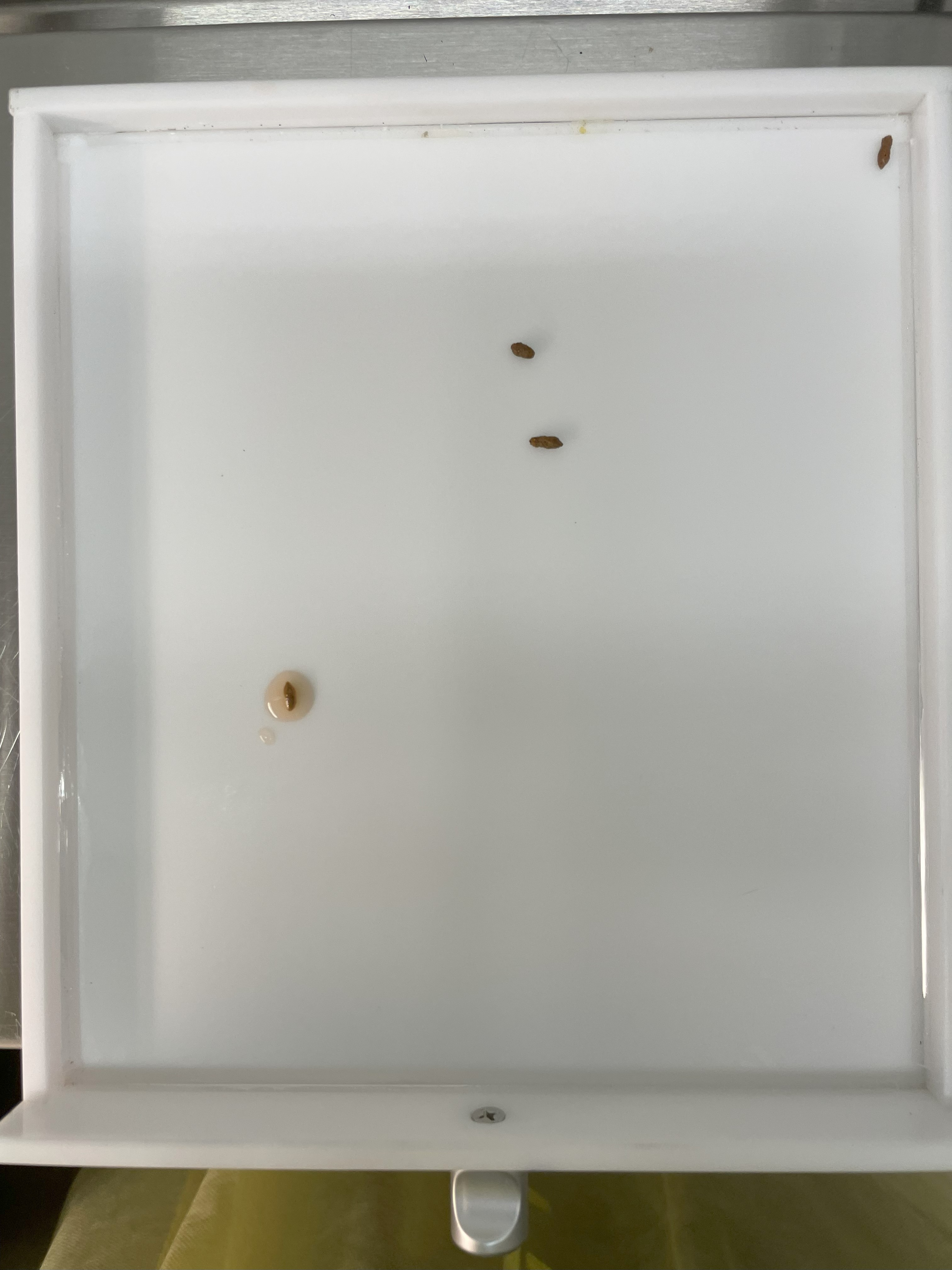

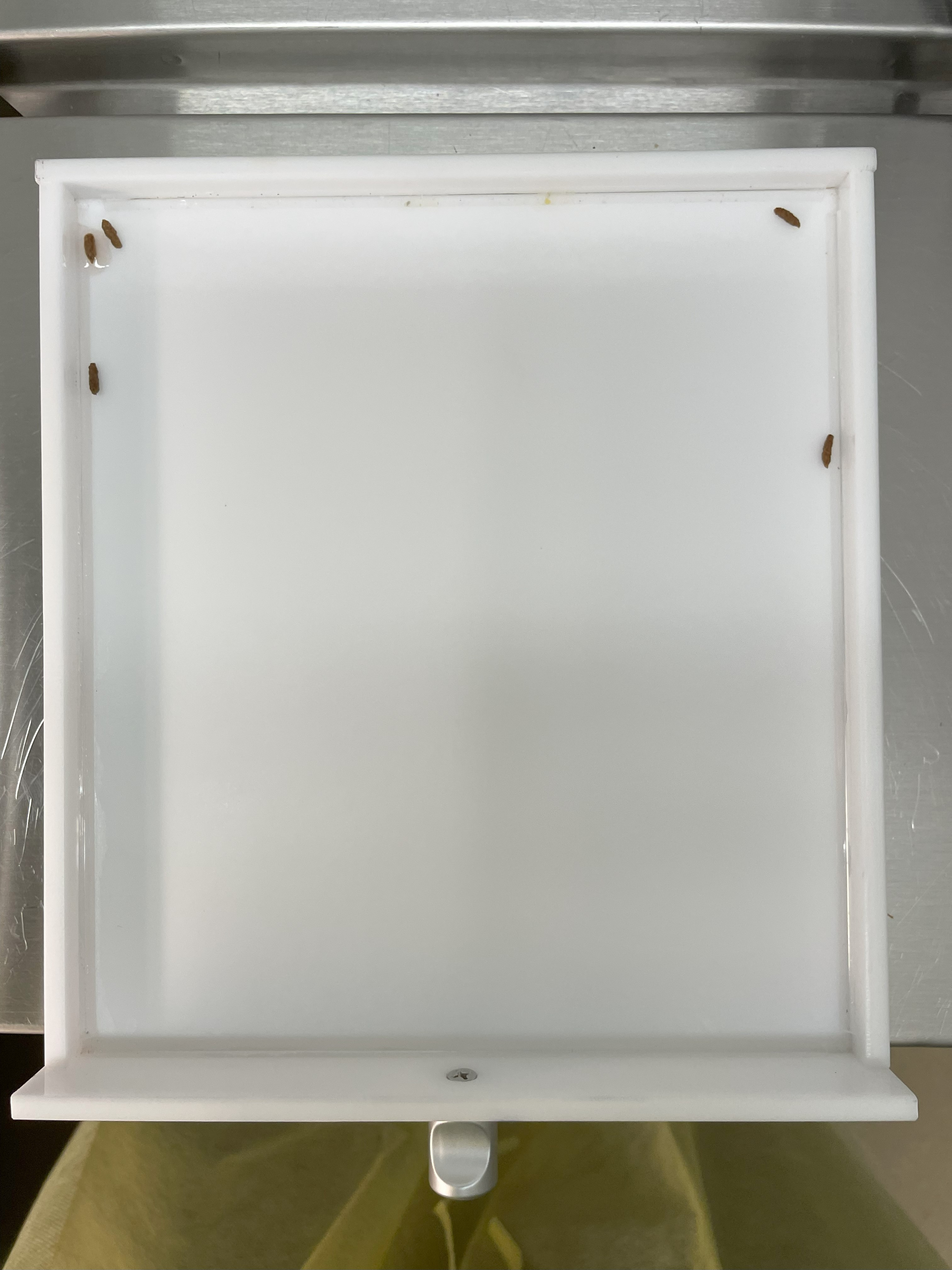

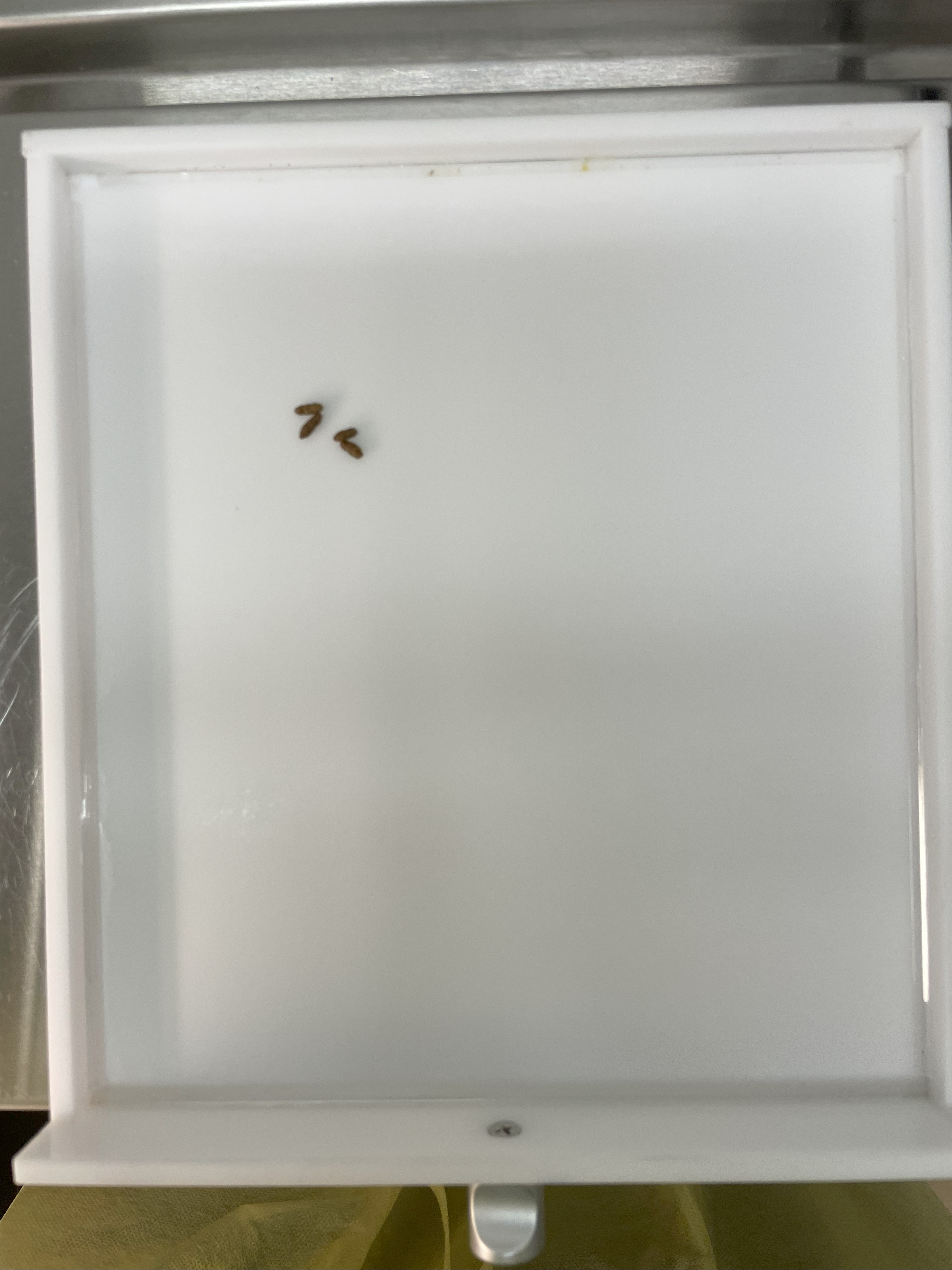

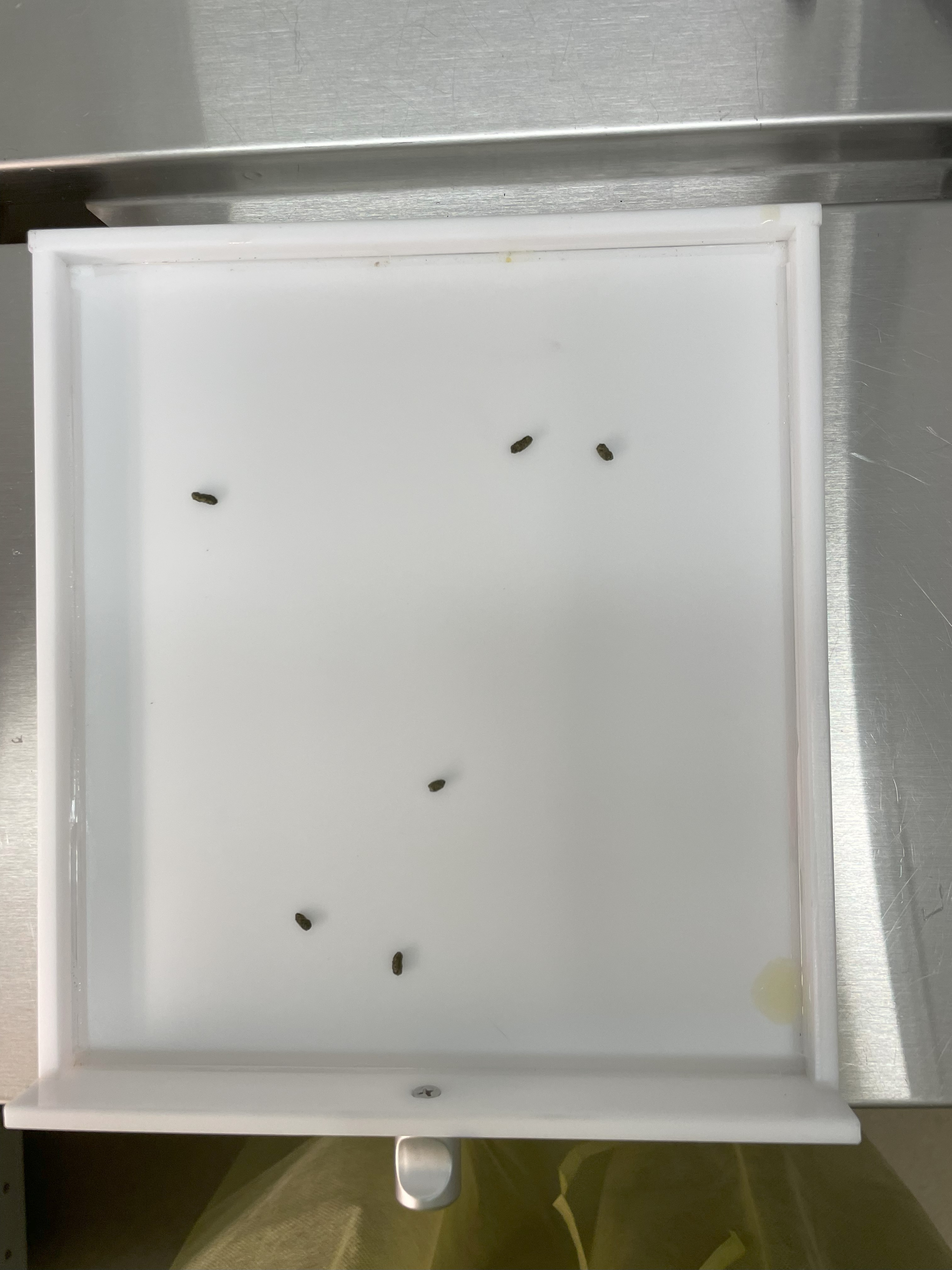

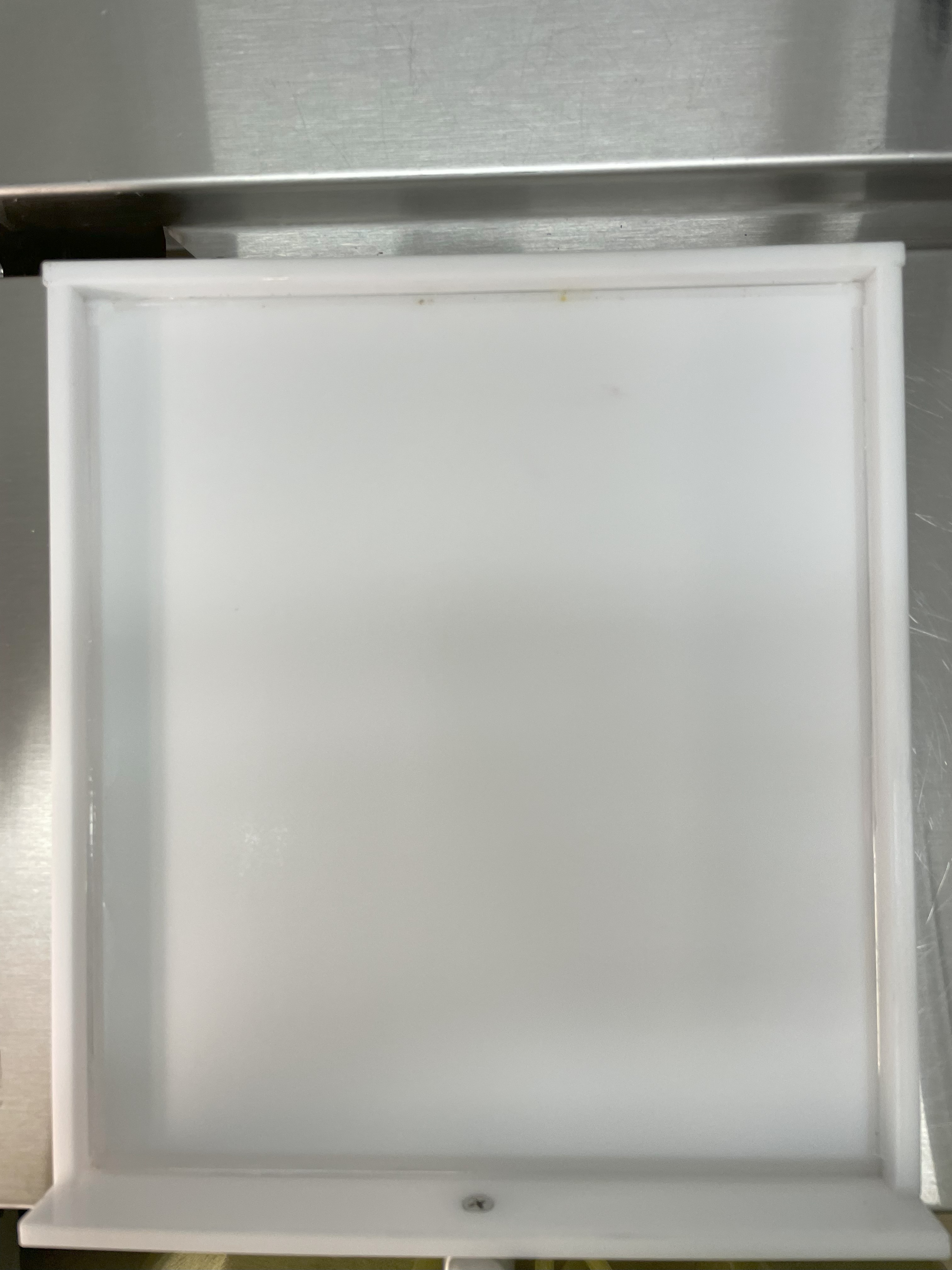

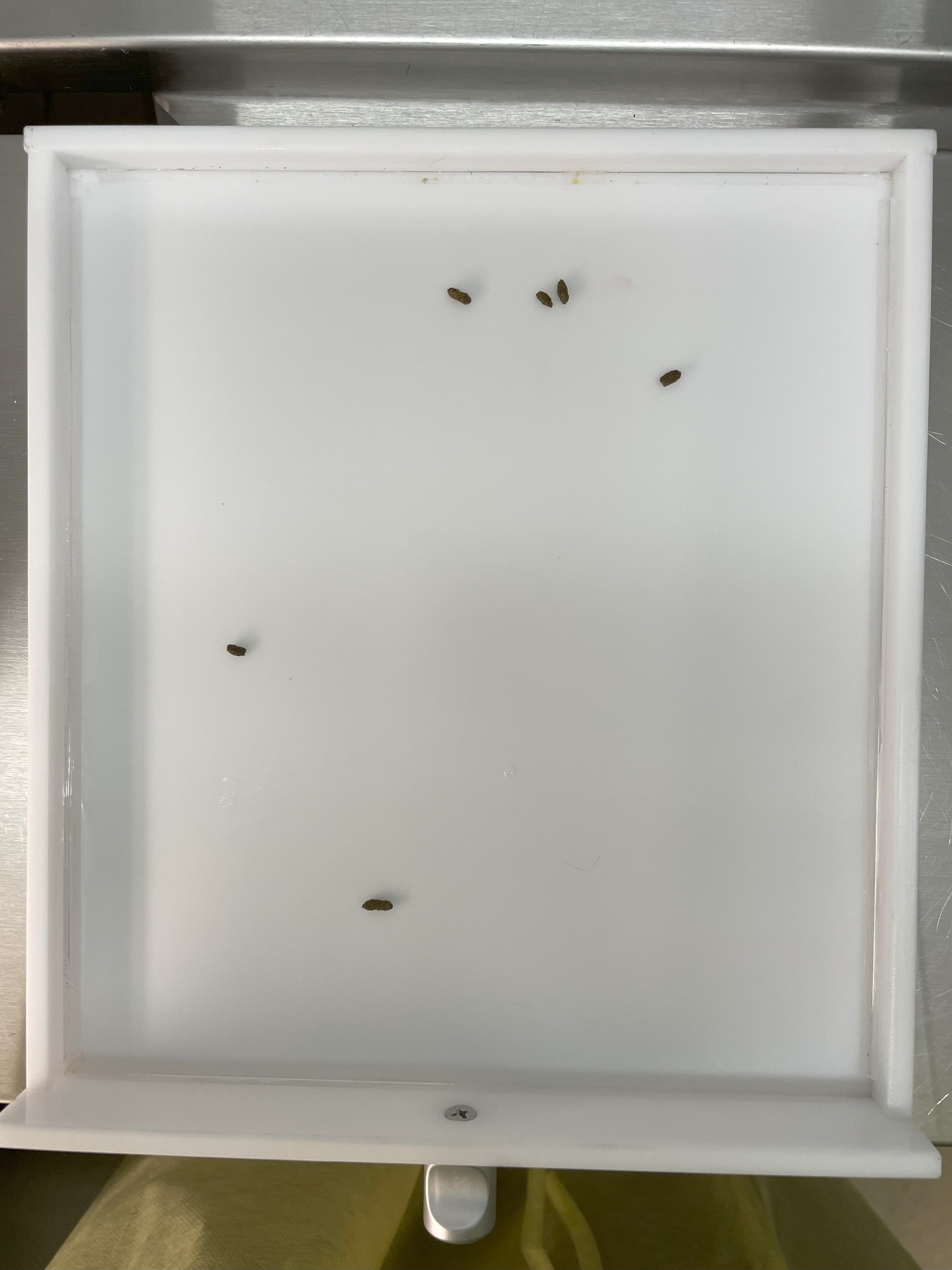

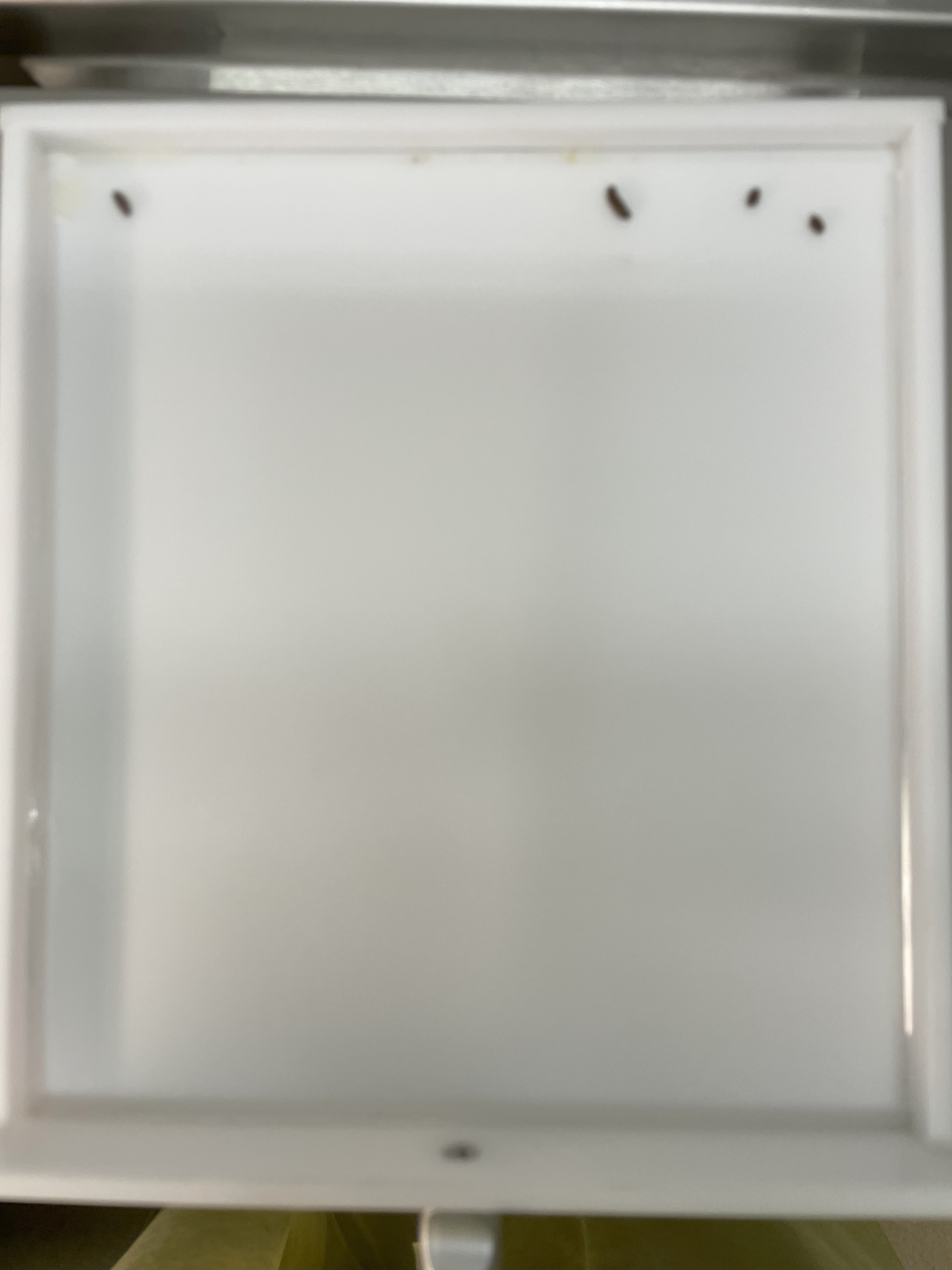

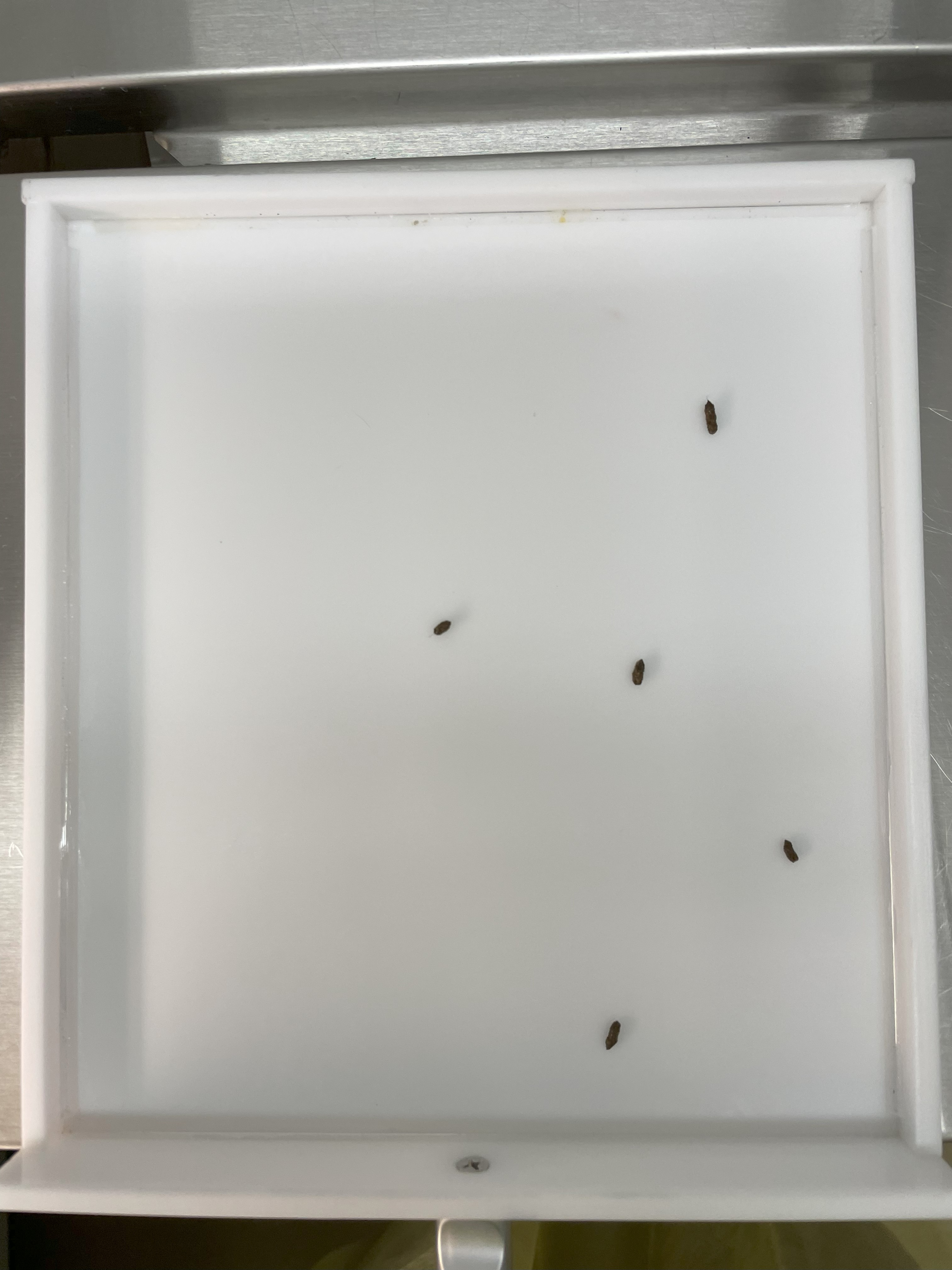

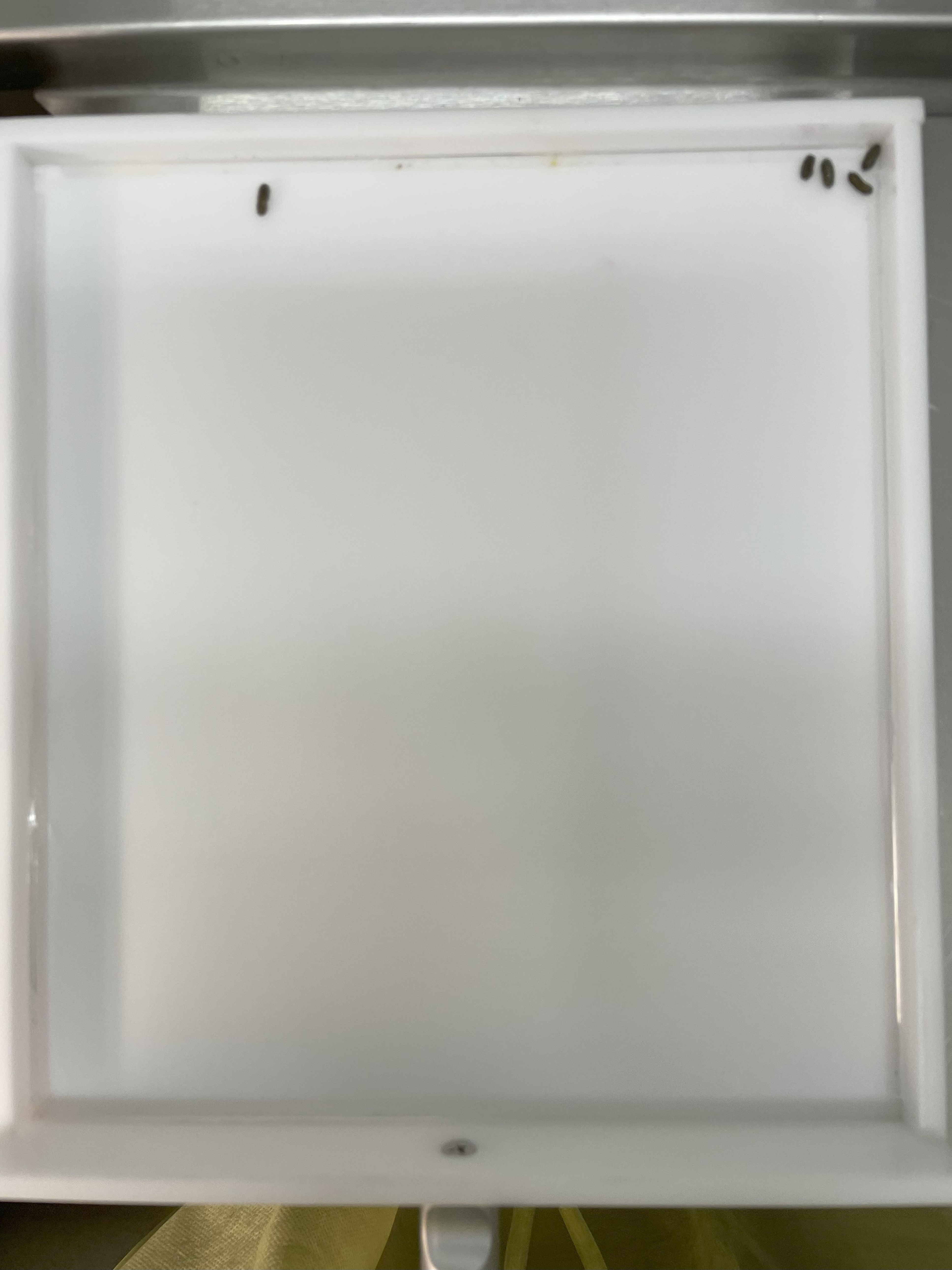

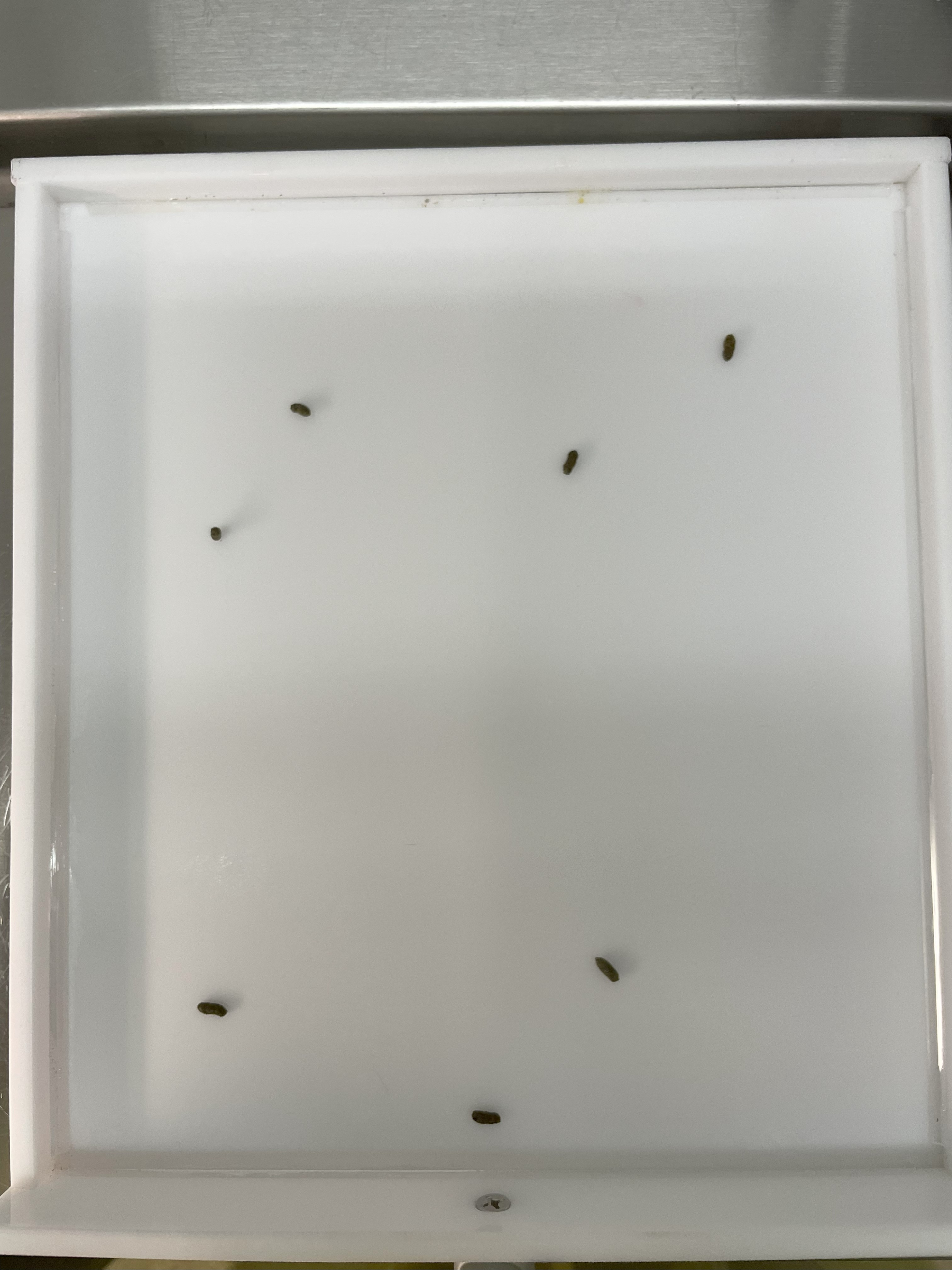

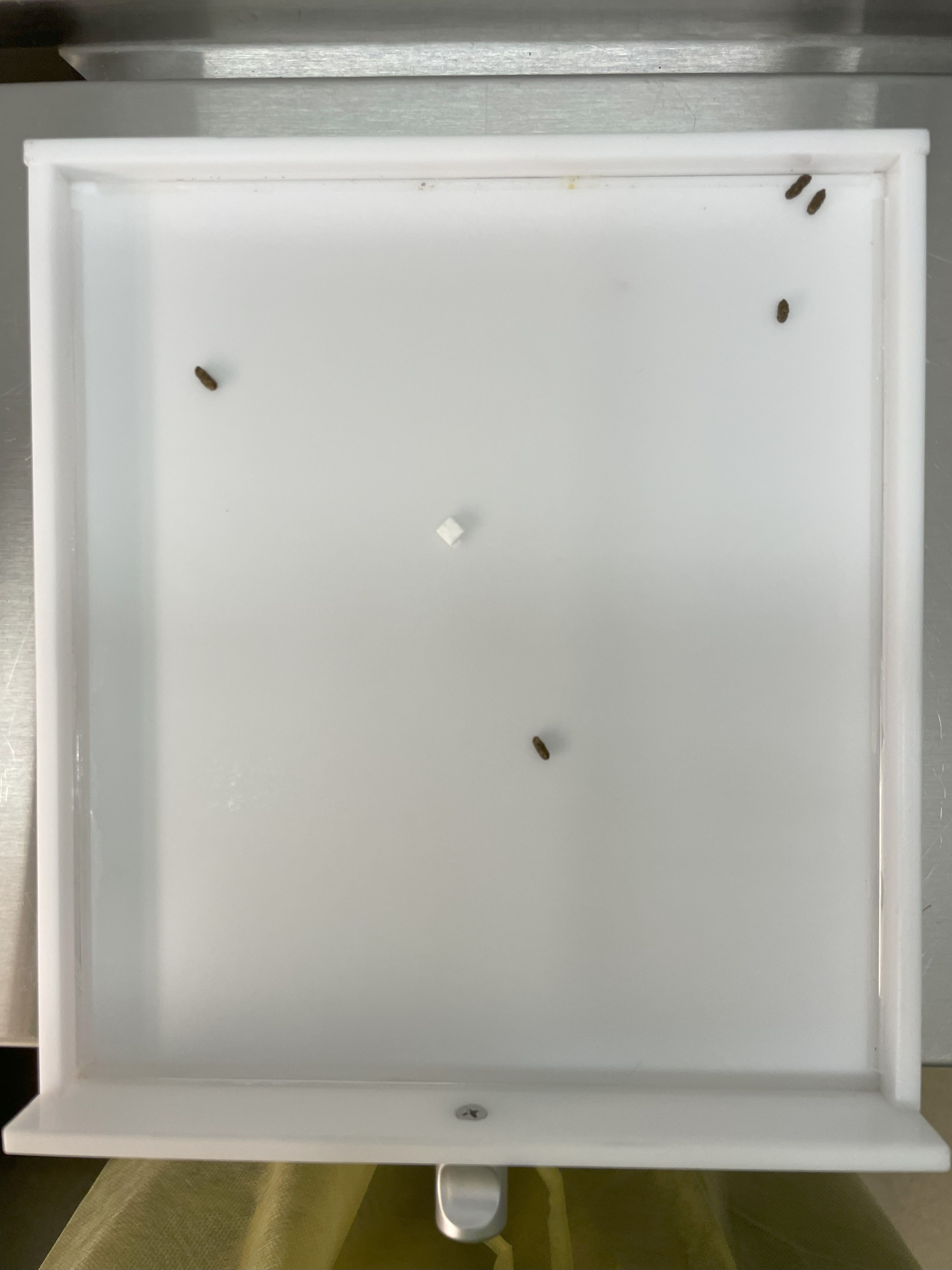

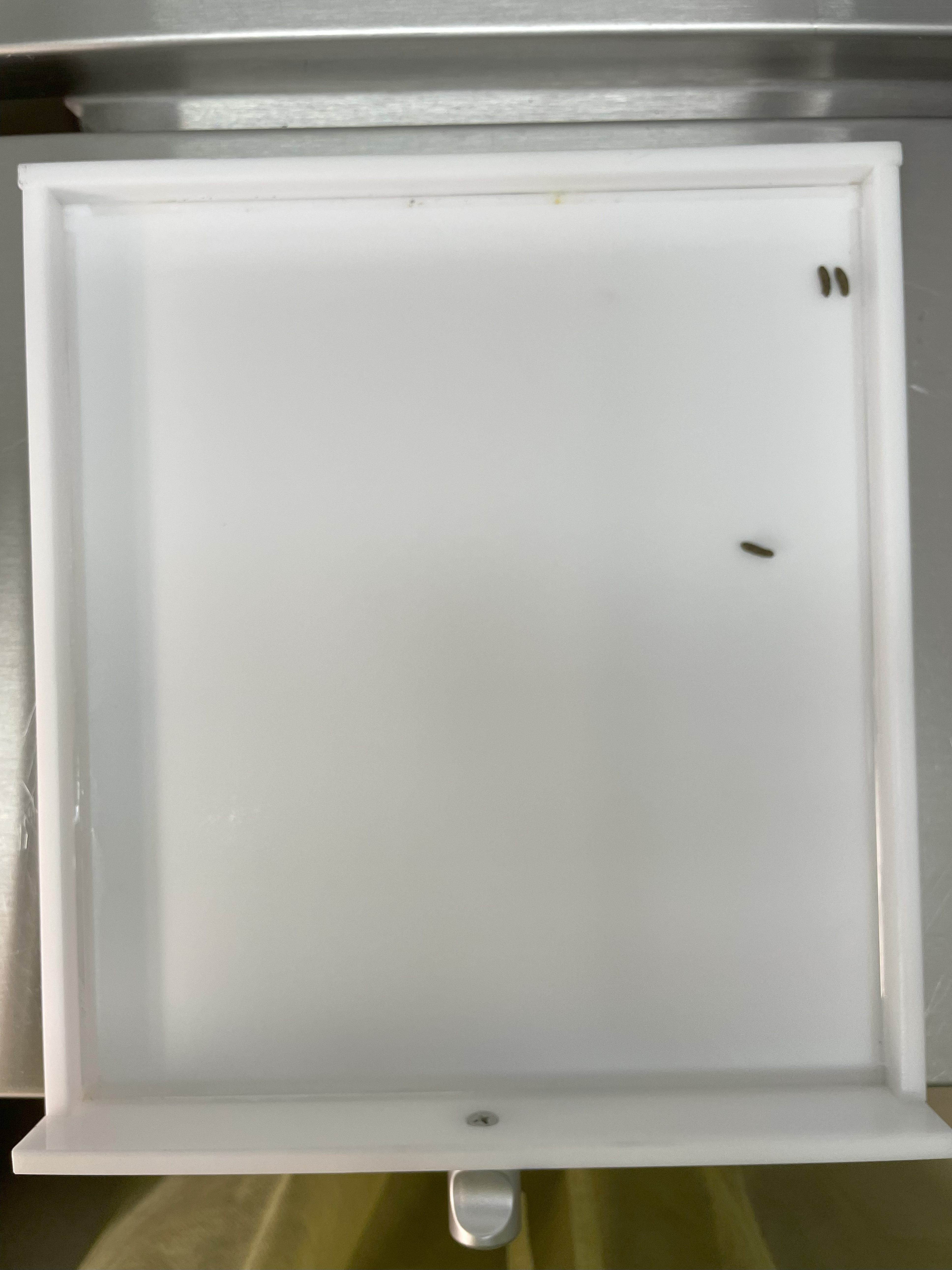

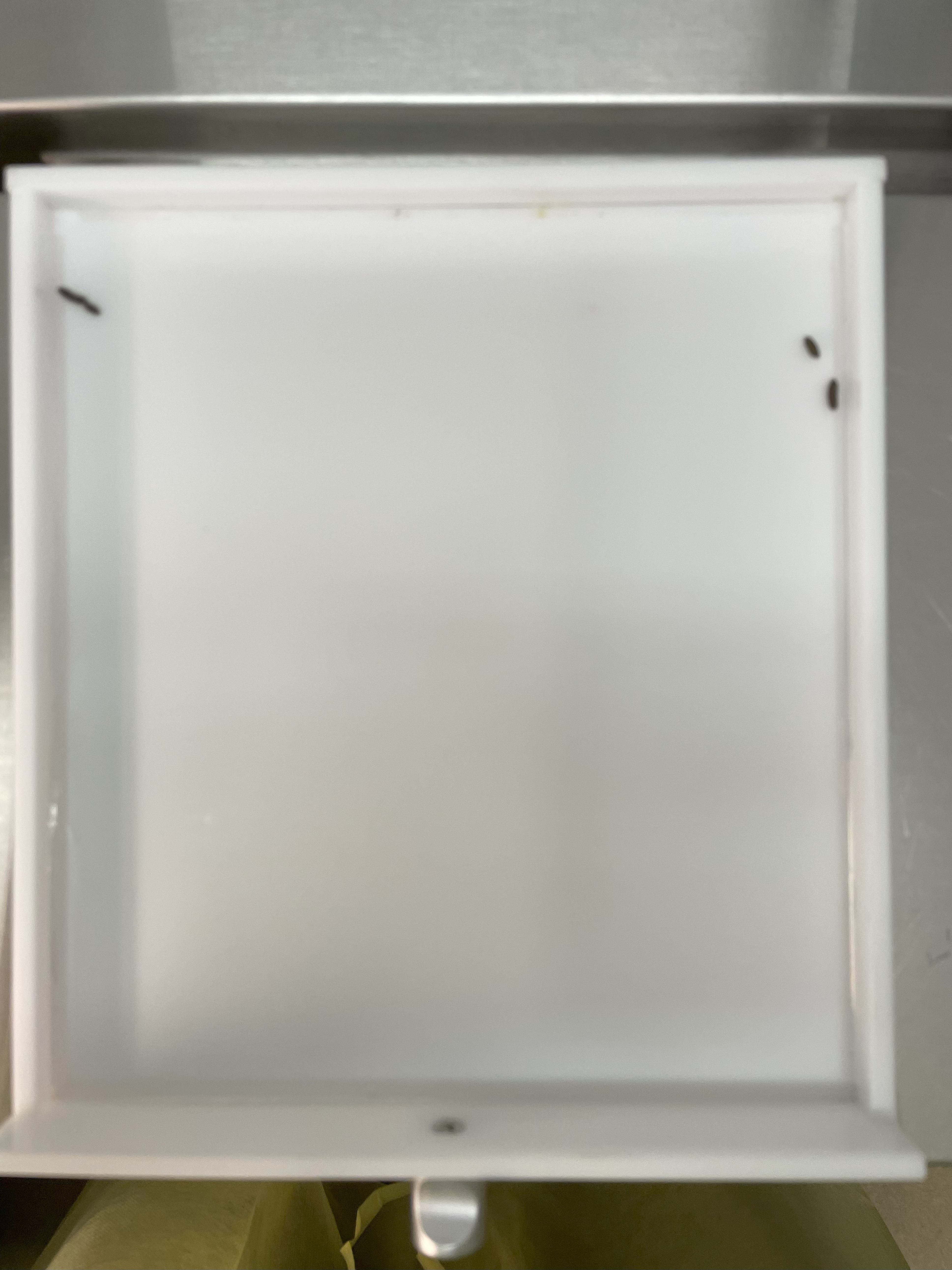
