## Supplementary Figure Caption for "Preliminary pharmacokinetics and *in vivo* studies indicate analgesic and stress mitigation effects of a novel NMDA receptor modulator"

Journal title: The Journal of Pharmacology and Experimental Therapeutics
Manuscript number: JPET-D-24-00077R1

**Supplementary Figure 1.** Predicted chemical structures and molecular weight of CNS4 metabolites 1 and 2.

**Supplementary Figure (video)2. CNS4 reduces electric shock sensation in male and female mice.** This video file shows four groups of C57BL mice (control-male, CNS4-male, control-female, CNS4-female) undergoing 2s of 0.5mA current shock after the following condition stimuli: 3min of acclimatization → 26s of 4kHz, 50% volume tone → 2s of white noise → 2s 0.5mA shock →17s gap. This was repeated 3X. Response to the first shock is shown in this video. S=seconds. CNS4, 100mg/kg, IP, treated 15-30min before shock, n=10 mice per group. The video file is uploaded as .mp4 file.

**Supplementary Figure 3. Effect of CNS4 on electric shock-induced fear conditioning (FC) on male and female mice.** Freezing episode (**A, D & G**), duration (**B, E&H**), latency (**C, F&I**), data collected on testing day and were compared between saline and CNS4 treated mice after tone1 (**A-C**), tone 2 (**D-F**) and tone 3 (**G-I**). Two-way ANOVA comparison is marked as a straight line on top of the bars, and post hoc tests are marked as downward pointing edges. Data points represent the biological replicates, n=10 per group. Data presented as mean ± sem. ns= not significant, $p>0.05$; *=$p<0.05$.

**Supplementary Figure 4.** Pictures show the fecal pellets observed on FC testing day (day 2). Pictures are arranged in the following order: Control male 1-5, control female 1-5, CNS4 male 1-5, and CNS4 female 1-5. Figure uploaded as a single .pdf file. Figure 4a & b contains pictures from the first and second set of experiments, respectively. There are twenty pictures in each set.
